# lncRNA *DIGIT* and BRD3 protein form phase-separated condensates to regulate endoderm differentiation

**DOI:** 10.1101/547513

**Authors:** Kaveh Daneshvar, M. Behfar Ardehali, Isaac A. Klein, Arcadia J. Kratkiewicz, Chan Zhou, Amin Mahpour, Brett M. Cook, Wenyang Li, Joshua V. Pondick, Sean P. Moran, Richard A. Young, Robert E. Kingston, Alan C. Mullen

## Abstract

Gene programs that control differentiation are regulated through the interplay between DNA, RNA, and protein. Cooperation among these fundamental cellular components can occur through highly structured interactions connecting domains formed by specific sequences of nucleotides, ribonucleotides, and/or amino acids and also through the assembly of biomolecular condensates. Here, we show that endoderm differentiation is regulated through the interaction of the long noncoding (lnc) RNA *DIGIT* and the bromodomain and extra-terminal (BET) domain family protein BRD3. BRD3 forms phase-separated condensates that contain *DIGIT*, occupies enhancers of endoderm transcription factors, and is required for endoderm differentiation. Purified BRD3 binds to acetylated histone H3 lysine 18 (H3K18ac) *in vitro* and occupies regions of the genome enriched in H3K18ac during endoderm differentiation, including the key transcription factors that regulate endoderm differentiation. *DIGIT* is also enriched in regions of H3K18ac, and depletion of *DIGIT* results in decreased recruitment of BRD3 to these regions. Our findings support a model where cooperation between *DIGIT* and BRD3 at regions of H3K18ac regulates the transcription factors that drive endoderm differentiation and suggest a broader role for protein-lncRNA phase-separated condensates as regulators of transcription in development.

## Introduction

Gene expression programs that determine cell identity are regulated by specific interactions between DNA, RNA, and protein, and these expression programs change during differentiation as new interactions are created and previous interactions fall apart. Signaling pathways frequently initiate differentiation (Basson, 2012), and activin or Nodal signaling provide the primary stimulus to direct human embryonic stem cells (hESCs) to differentiate into definitive endoderm (D’Amour et al., 2005; Kubo et al., 2004). Downstream of signaling pathways, differentiation is regulated genetically and epigenetically by coding and noncoding elements of the genome (Gifford et al., 2013). Long noncoding RNAs (lncRNAs) are components of the regulatory circuits that control pluripotency and differentiation into each germ layer (Flynn and Chang, 2014), and the lncRNAs *DIGIT, DEANR1*, and *EVX1* promote mesendoderm and/or definitive endoderm differentiation (Bell et al., 2016; Daneshvar et al., 2016; Jiang et al., 2015; Luo et al., 2016).

*DIGIT* is an lncRNA that regulates differentiation of human and murine ESCs into definitive endoderm (Daneshvar et al., 2016). *DIGIT* controls endoderm differentiation, at least in part, through regulation of the transcription factor Goosecoid (GSC). *DIGIT* is divergently transcribed from the gene encoding GSC, but *DIGIT* does not need to be expressed in close proximity to *GSC* in the genome to regulate *GSC* expression. The mechanism by which *DIGIT* regulates *GSC* and controls endoderm differentiation is unknown. Therefore, we employed an RNA-centric approach to define the protein interactome of *DIGIT* and found that BRD3, a member of the bromodomain and extra-terminal (BET) domain family of proteins, showed the strongest interaction.

The BET family of proteins is composed of BRD2, BRD3, BRD4, and testis-specific BRDT (Paillisson et al., 2007). BET proteins contain two bromodomain motifs, which bind acetylated histones (Filippakopoulos et al., 2012; Kanno et al., 2004) and an extra-terminal domain that enables them to interact with other proteins including transcription factors, transcription co-activators, pause release factors, and Mediator proteins (Deeney et al., 2016; Jang et al., 2005; Paillisson et al., 2007; Wai et al., 2018). BRD4 has also been the focus of many studies due to its role as a regulator of MYC in cancer (Delmore et al., 2011; Zuber et al., 2011).

BRD2, BRD3, and BRD4 are ubiquitously expressed, but unique functions have been attributed to specific family members in development. Loss of either BRD4 or BRD2 leads to embryonic lethality, but death occurs at different stages, with *Brd4*^−/−^ embryos dying before implantation (Houzelstein et al., 2002) and *Brd2*^−/−^ embryos dying at mid-gestation between E9.5 and E11.5 (Shang et al., 2009). BRD4, but not BRD2 or BRD3, also regulates pluripotency and self-renewal of murine ESCs (Di Micco et al., 2014), while BRD2 promotes expression of Nodal when ESCs are released from pluripotency (Fernandez-Alonso et al., 2017). The different activities of BET family proteins in ESCs are associated with different patterns of genome occupancy, where BRD4 is enriched primarily at enhancers, and BRD2 and BRD3 are enriched at promoters (Engelen et al., 2015). Furthermore, BRD4 has been shown to form phase-separated droplets in murine ESCs, which have been proposed to promote recruitment of transcription machinery to regulate gene expression (Sabari et al., 2018). While critical roles for BRD2 and BRD4 have been established in early development, the function of BRD3 is not well understood.

Here we report that *DIGIT* interacts with BRD3 and is found with BRD3 in phase-separated condensates during endoderm differentiation. With differentiation, BRD3 occupies new sites at endoderm genes, many of which are also regulated by *DIGIT*. Recombinant BRD3 preferentially recognizes histone H3 lysine 18 acetylation (H3K18ac), a modification produced by the transcription co-activators CBP/p300 (Jin et al., 2011), and BRD3 co-occupies the genome with H3K18ac. *DIGIT* is also enriched at sites of H3K18ac, and depletion of *DIGIT* reduces BRD3 occupancy at these sites. We propose that BRD3 is recruited to sites of H3K18ac to form phase-separated condensates and promote transcription, and this recruitment of BRD3 is dependent on *DIGIT*. Thus, the interaction between *DIGIT* and BRD3 at sites of H3K18ac leads to activation of genes that regulate endoderm differentiation.

## Results

### Design of an aptamer-based approach to define the DIGIT-protein interactome

*DIGIT* transcripts are retained in the nucleus and regulate genes that control endoderm differentiation (Daneshvar et al., 2016). We asked whether *DIGIT* transcripts directly interact with nuclear proteins in hESCs undergoing endoderm differentiation. We fused the 3’ end of the cDNA encoding *DIGIT* to four copies of an aptamer with high affinity for streptavidin (4xS1m) (Leppek and Stoecklin, 2014; Srisawat and Engelke, 2001). We performed single-molecule RNA-FISH (smFISH) in hESCs after transient transfection with a plasmid expressing *DIGIT-4xS1m* and differentiated hESCs toward definitive endoderm. Under these conditions, the *DIGIT-4xS1m* chimeric RNA molecules are retained in the nucleus (Figure 1A), as previously described for endogenous *DIGIT*.

**Figure 1.**
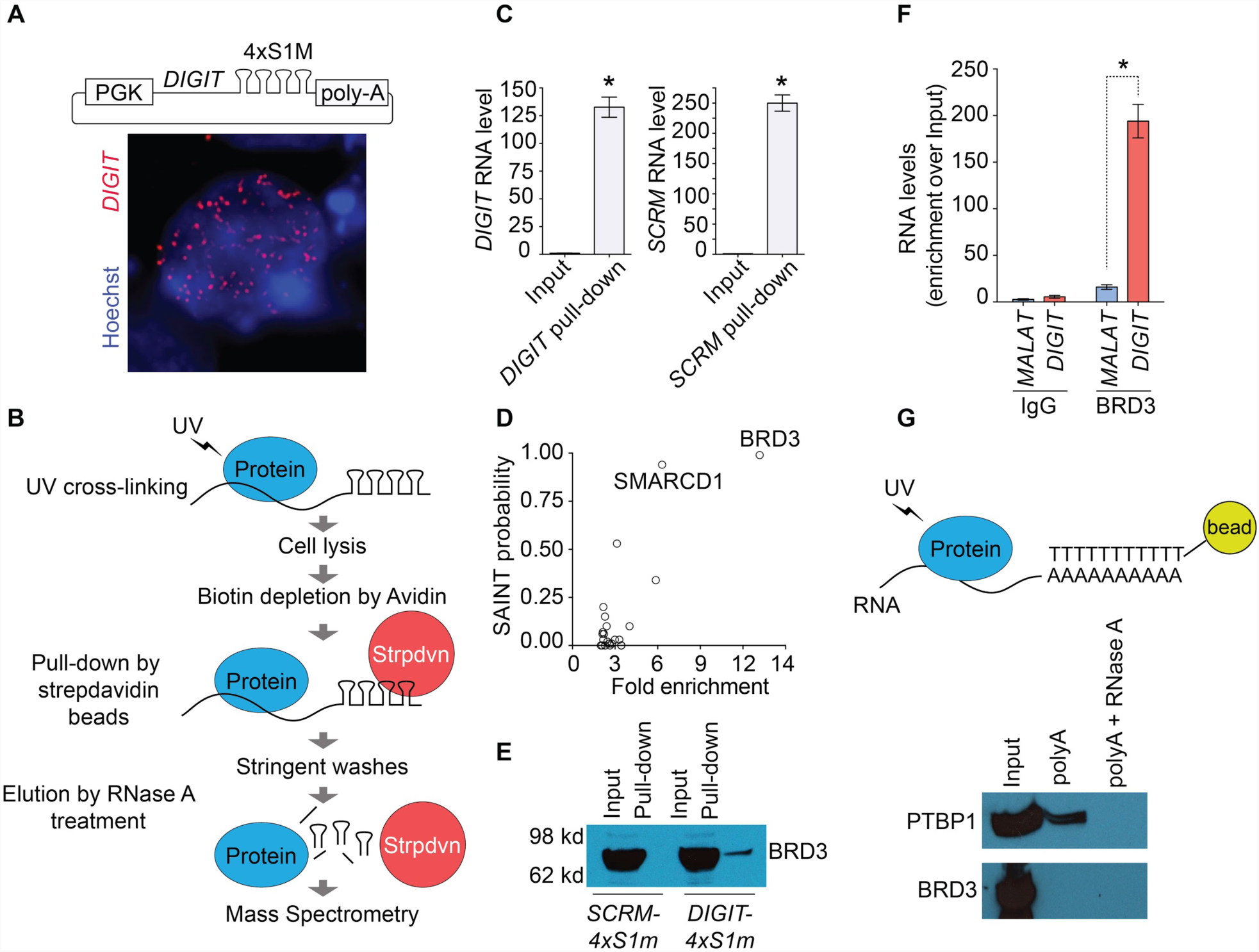
*DIGIT* interacts with BRD3 in definitive endoderm cells. **(A)** Schematic map of the construct expressing *DIGIT* fused to four S1m aptamers (*DIGIT-4xS1m*, top). RNA-FISH was performed to detect ectopically expressed *DIGIT-4xS1m* chimeric RNA in hESCs (bottom). *DIGIT* transcripts are red and the nucleus is shown in blue (Hoechst). **(B)** Strategy for UV cross-linking, streptavidin-assisted pull-down of ectopically-expressed *DIGIT*, and mass spectrometry analysis of interacting proteins in endoderm cells. **(C)** qRT-PCR was performed to quantify enrichment of *DIGIT-4xS1m* (left) and scrambled *DIGIT* (*SCRM-4xS1m*) (right) after streptavidin pull-down. * indicates p <0.0001. **(D)** Proteomic analysis of the mass spectrometry data for *DIGIT-4xS1m* compared to *SCRM-4xS1m* and to *DIGIT* without aptamers on day 4 of endoderm differentiation. The x-axis indicates fold enrichment compared to controls. The y-axis indicates the probability score for each interaction based on the SAINT (Significance Analysis of INTeractome) score. **(E)** Immunoblot for BRD3 was performed using cell lysates on day 4 of endoderm differentiation (5% input) and then after streptavidin-assisted pull-down of *SCRM-4xS1m* and *DIGIT-4xS1m*. **(F)** RNA-Immunoprecipitation (RIP) was performed on day 4 of endoderm differentiation using an anti-BRD3 antibody or an IgG isotype control. qRT-PCR was performed to quantify expression of *DIGIT*. Enrichment of lncRNA *MALAT1* is shown as a negative control. * indicates p <0.0001. **(G)** hESCs were differentiated for 4 days before UV cross-linking and pull-down of polyadenylated (polyA) RNAs. The immunoblot was performed to detect protein levels of PTBP1 (top) and BRD3 (bottom). PolyA selection after RNase A treatment was performed to confirm that enrichment was due to interaction with polyA RNA.

We next asked if *DIGIT-4xS1m* chimeric transcripts were precipitated with streptavidin beads in UV cross-linked cells (Figure 1B). We tested *DIGIT* enrichment with increasing concentrations of potassium chloride (KCl) in the wash buffer and determined that at 350 mM KCl *DIGIT* enrichment is well above *GAPDH* (Figure S1A). qRT-PCR showed 125-fold enrichment of *DIGIT-4xS1m* compared to *GAPDH* mRNA after streptavidin precipitation (Figure 1C). Together, these experiments show that the 4xS1m aptamer can be fused to lncRNAs and used to enrichm of lncRNAs in UV cross-linked cells.

### DIGIT interacts with BRD3

To identify the proteins that interact with *DIGIT* transcripts in differentiating cells, we transiently expressed *DIGIT-4xS1m* transcripts in hESCs that were differentiating toward endoderm. As a control, we ectopically expressed a scrambled form of *DIGIT* fused to 4xS1m (*SCRM-4xS1m*) and the *DIGIT* transcript without the 4xS1m aptamers. We performed UV cross-linking followed by streptavidin precipitation in cells expressing each transcript. Proteins released by RNase A digestion were resolved on a polyacrylamide gel prior to analysis by mass spectrometry (MS). After obtaining the spectral counts for each sample, we used a proteomics analysis pipeline to identify candidate proteins that interact with *DIGIT* (Mellacheruvu et al., 2013) (Crapome.org). We set a threshold of >90% confidence and greater than four-fold enrichment over controls. This analysis identified BRD3 as the protein with the highest interaction score (Figure 1D and 1E, Table S1). The other protein to pass this threshold was SMARCD1, a component of the SWI/SNF chromatin remodeling complex (Oh et al., 2008) which is involved in embryonic stem cell differentiation (Alajem et al., 2015).

Since aptamer experiments were performed using ectopically expressed *DIGIT* transcripts that localized to the nucleus, it is possible that differences in the number of *DIGIT* molecules present in a cell during endoderm differentiation could lead to interactions that are not formed with endogenous *DIGIT*. To validate the interaction of BRD3 with endogenous *DIGIT* during endoderm differentiation, we performed RNA-immunoprecipitation (RIP) in nuclear lysates of differentiating cells using an antibody against BRD3 and quantified enrichment of the *DIGIT* transcript (Figure 1F). RNA precipitated with BRD3 demonstrated a 175-fold increase in *DIGIT* compared to RNA precipitated with IgG. This enrichment was observed at a much lower level for *MALAT1*, an abundant nuclear lncRNA. These results suggest that BRD3 interacts with *DIGIT* during endoderm differentiation.

### BRD3 does not broadly associate with RNA and shows specificity for DIGIT compared to BRD2 and BRD4

BRD2, BRD3, and BRD4 have been shown to interact with a small number of RNAs transcribed from enhancer regions (eRNAs) (Rahnamoun et al., 2018). However, the specificity of the interactions and whether these proteins bind to other RNA species are not known. *DIGIT* transcripts are polyadenylated (polyA) (Sigova et al., 2013), and we asked if BRD3 associates broadly with polyA RNAs. We UV cross-linked hESCs and precipitated polyA RNAs from nuclear extracts using oligo-dT-coated beads (Castello et al., 2016). As a positive control, we performed an immunoblot for Polypyrimidine Tract Binding Protein 1 (PTBP1), a protein that is associated with RNA splicing (Garcia-Blanco et al., 1989) and has been shown to interact with lncRNAs (Lin et al., 2014; Ramos et al., 2015) (Figure 1G). Despite enrichment of PTBP1 with polyA RNA precipitation, BRD3 was not detected in association with polyA RNAs. Beta-ACTIN served as a negative control for this experiment, as we did not find evidence in the literature of RNA-binding activity for this protein. Treatment with RNase A also confirmed that the interaction detected between PTBP1 and polyA RNAs was dependent on the presence of RNAs. These results suggest that BRD3 interacts with *DIGIT*, but BRD3 does not broadly interact with polyA RNAs.

BRD2, BRD3, and BRD4 but not BRDT are expressed upon endoderm differentiation (Figure S1B), and we asked if the other BET family proteins present during differentiation also interact with *DIGIT*. We performed RIP to quantify the interaction between *DIGIT* and BRD2, BRD3, and BRD4 during endoderm differentiation. Consistent with our previous analyses, we observed enrichment of *DIGIT* with precipitation of BRD3. However, enrichment of *DIGIT* was significantly reduced with precipitation of BRD2 and BRD4 compared to BRD3 (Figure S1C). These findings reveal that BRD3 exhibits specificity in its interactions with RNA, and BRD3 has a higher affinity for *DIGIT* than either BRD2 or BRD4.

### DIGIT localizes within large BRD3 puncta

To further investigate the interactions between *DIGIT* and BRD3, we visualized *DIGIT* and BRD3 within the nuclei of differentiating cells. We observed that 18% of *DIGIT* transcripts overlap with BRD3 staining (Figure 2A and 2B). This overlap was significantly higher than the overlap observed between BRD3 protein and *GSC* mRNA (<1%) (Figure 2B, left). *DIGIT* is primarily retained in the nucleus while the majority of *GSC* mRNA is in the cytoplasm (Daneshvar et al., 2016). For this analysis, we focused only on transcripts localized to the nucleus for both *DIGIT* (379) and *GSC* mRNA (2412). The higher number of *GSC* transcripts reflects the larger number of *GSC* mRNA molecules in cells at day 4 of endoderm differentiation. The images also revealed that BRD3 is not evenly distributed through the nucleus. Instead, BRD3 tends to form puncta, a phenomenon that has been described for BRD4 (Sabari et al., 2018). We observed that, when compared to nuclear *GSC* mRNA molecules that overlap with BRD3 staining, *DIGIT* transcripts are more likely to localize in larger BRD3 puncta (Figure 2B, right). The analysis also revealed multiple sites of overlap between BRD3 and *DIGIT* in nuclei, indicating that we are not simply observing sites of *DIGIT* transcription inside BRD3 condensates as could be hypothesized if there were only 1-2 sites of overlap in each cell. These results suggest that *DIGIT* transcripts frequently associate with BRD3 puncta during endoderm differentiation.

**Figure 2.**
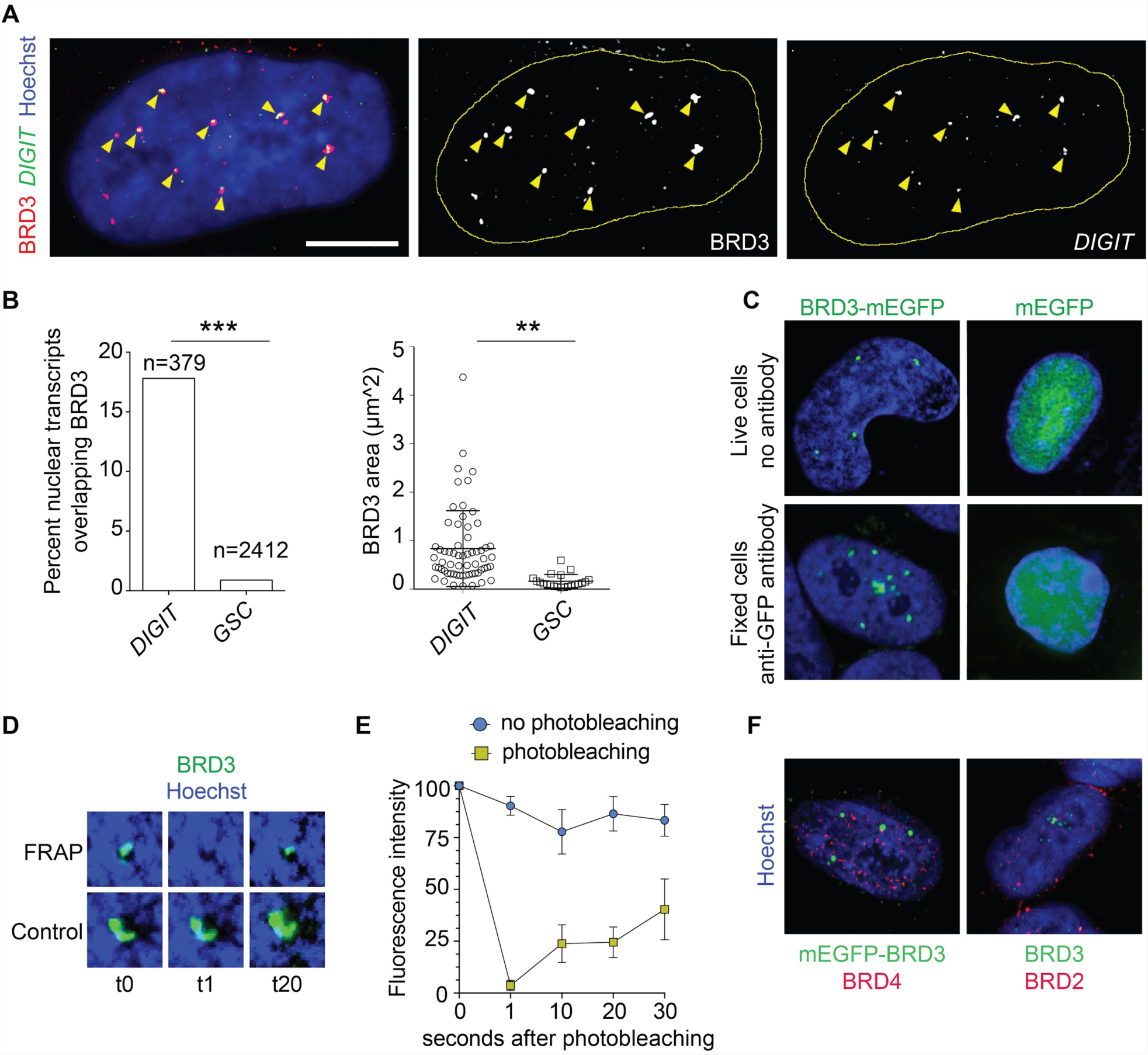
BRD3 forms protein-RNA phase-separated condensates. **(A)** Sequential immunofluorescence and RNA-FISH shows the distribution of BRD3 protein (red) and *DIGIT* RNA (green) within the nucleus (blue) on day 4 of endoderm differentiation (first panel). Areas of overlap are shown in yellow and marked with yellow arrows. The second panel shows BRD3 protein (white) and the third panel shows *DIGIT* RNA (white) for the same cell. The nuclei are outlined in yellow for the second and third panel. Scale bar represents 10 μm. **(B)** The bar chart shows the number of *DIGIT* lncRNA and *GSC* mRNA molecules inside the BRD3 staining area (left). *** indicates p <0.0001 (Fisher exact test). The plot on the right shows the distribution of *DIGIT* and *GSC* transcripts that overlap with BRD3, as a function of BRD3 condensate size. ** indicates p <0.001. **(C)** Live (top) and fixed cell (bottom) fluorescence images of BRD3 condensates (green) are shown in nuclei (blue). **(D)** FRAP analysis of BRD3 condensates (green) before photobleaching (t0) and 1 second (t1) and 20 seconds (t20) after photobleaching. **(E)** Quantification of the FRAP experiment imaged in (D). **(F)** Co-immunostaining of BRD3 (green) with BRD4 (red, left) or BRD2 (red, right) in endoderm cells.

### BRD3 shows properties of liquid-liquid phase-separated condensates

Disordered proteins can form biological condensates and interact with RNA (Castello et al., 2016; He et al., 2016). Analysis of the primary sequence of BRD3 using PONDR VSL2 showed that 75% of the long isoform of BRD3 and 67% of the short isoform of BRD3 are disordered (Figure S2A). To further investigate the possibility that BRD3 puncta are phase-separated condensates, we first asked if BRD3, expressed as a fusion protein with EGFP, forms condensates in live cells. We used the CRISPR/Cas system to insert a monomeric EGFP (mEGFP) gene into the 5’ end of the endogenous *BRD3* gene (Figure S2B and S2C). The generation of fused mEGFP-BRD3 was confirmed by immunoblot after pull-down with an antibody recognizing GFP (Figure S2D). Confocal microscopy of differentiating cells confirmed the formation of BRD3 puncta in both live and fixed endoderm cells (Figure 2C). As a control, we generated hESCs that stably express mEGFP fused to a nuclear localization signal (NLS). Live imaging and immunostaining of GFP in these mEGFP-expressing cells show that mEGFP by itself is not capable of forming puncta in differentiating cells.

Phase-separated condensates have a liquid-like state and can exchange with their surroundings (Boija et al., 2018; Lu et al., 2018; Sabari et al., 2018). To test whether BRD3 puncta display rapid exchange kinetics with their surroundings, we performed Fluorescence Recovery After Photobleaching (FRAP) analysis of EGFP-tagged BRD3 proteins in live endoderm cells. After photobleaching, the BRD3 puncta are able to recover up to 20% in 10 seconds and up to 38% in 30 seconds (Figure 2D and 2E). These results showed that BRD3 molecules can form puncta with properties consistent with liquid-liquid phase-separated condensates, and that components of these condensates can dynamically exchange molecules with their surroundings.

The evidence from imaging suggests that BRD3 forms phase-separated condensates in hESCs and with endoderm differentiation (Figure 2C, 2D, and S2E). BRD4 forms condensates in hESCs (Sabari et al. 2018), and BRD2, BRD3, and BRD4 all have large domains of predicted disorder (Figure S2A). We found that all three proteins form puncta with endoderm differentiation and that BRD3 forms puncta that are separate from BRD4 and BRD2 (Figure 2F). Overall, these findings suggest that each BET family protein form distinct puncta with endoderm differentiation.

### Formation of BRD3 condensates is concentration-dependent and can enrich RNA

The proteins that compose phase-separated biomolecular condensates may form liquid droplets *in vitro* (Banani et al., 2017). To test the protein concentration dependency of phase-separated BRD3 condensates, we produced recombinant BRD3 fused to mEGFP and purified this protein from eukaryotic cells (Figure S3A). A titration of the protein showed that BRD3 can form droplets at concentrations as low as 5 μM (Figure 3A). Phase-separated droplets which rely on electrostatic interactions are sensitive to salt, so we asked if recombinant BRD3 can form droplets at physiological salt concentrations. A titration of the salt showed that BRD3 can form droplets in salt concentrations as high as 300 mM (Figure 3B). We next asked if BRD3 condensates formed *in vitro* interact with the *DIGIT* transcript. We synthesized Cy3-labeled *DIGIT* (Figure S3B) and induced *in vitro* formation of BRD3 condensates in the presence of Cy3-*DIGIT*. We observed that the Cy3-*DIGIT* molecules were concentrated within BRD3 condensates (Figure 3C), while Cy3-*DIGIT* molecules alone remain dispersed (Figure 3C, far right). These results show that BRD3 forms phase-separated condensates that recruit *DIGIT in vitro*. Although these results do not provide concrete evidence that BRD3 condensates have specific affinity for the *DIGIT* transcript, they support a model where BRD3 condensates form a suitable environment for interaction with RNAs. Additional studies will be needed to address the selectivity of BRD3 condensates for specific RNA species.

**Figure 3.**
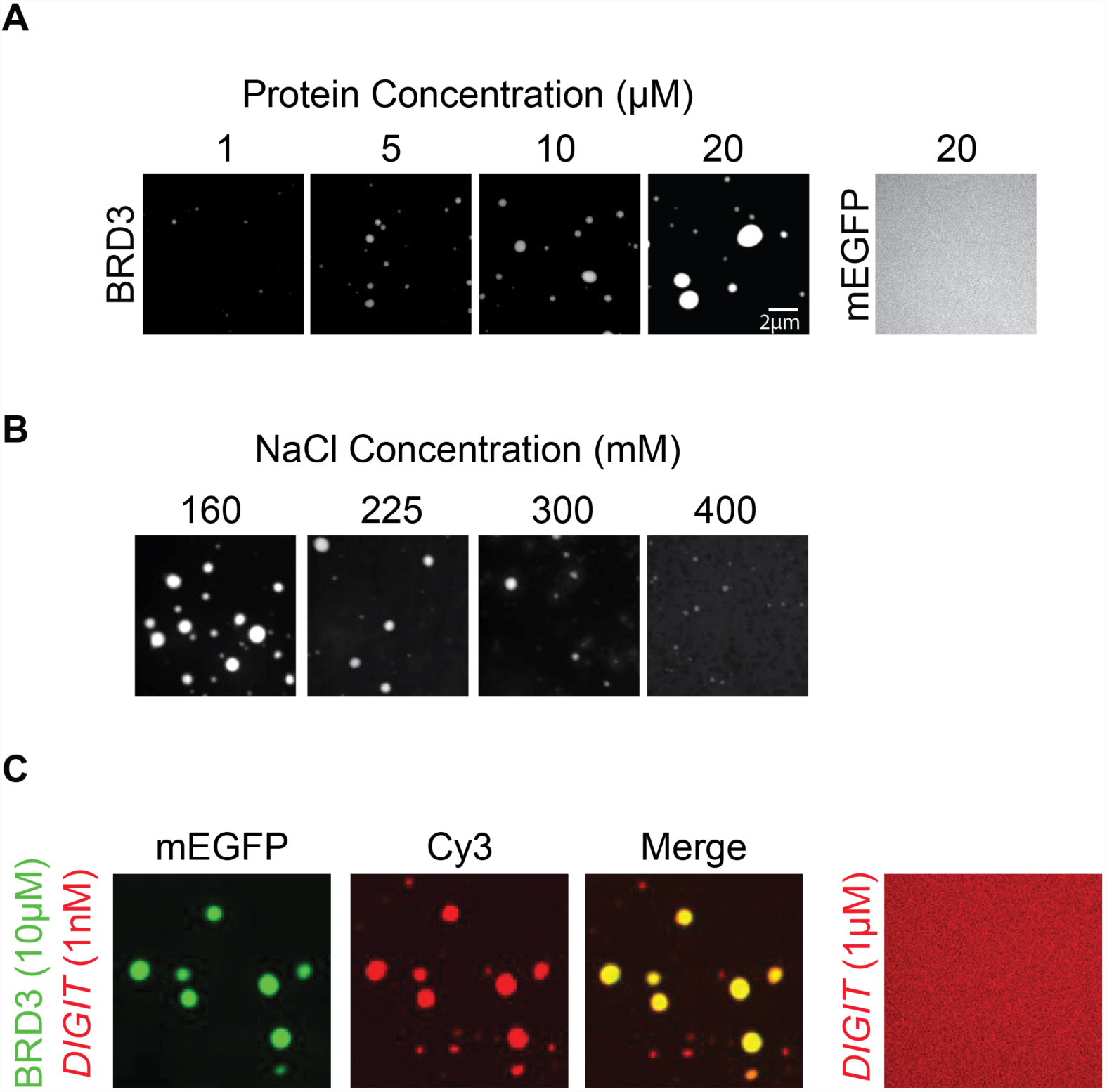
BRD3 protein forms phase-separated condensates *in vitro.* **(A)** Purified recombinant BRD3 fused to monomeric (m) EGFP forms phase-separated condensates, indicated by white circles on a black background, at the indicated concentrations. Purified recombinant mEGFP is used as control (far right). **(B)** *In vitro* formation of BRD3 condensates as a function of salt concentration. 10 μM BRD3-mEGFP was incubated at the indicated salt concentrations and assessed for droplet formation. **(C)** BRD3 condensates can enrich labeled *DIGIT*. Enrichment of fluorescently labeled *DIGIT* (Cy3, red) within BRD3-mEGFP droplets (green). The overlap between BRD3-mEGFP and *DIGIT*- Cy3 is shown in yellow (merge). Cy3-*DIGIT* alone is shown on the far right (1 μM). Concentrations of BRD3 and *DIGIT* used for mixing are indicated at the far left.

### BRD3 regulates definitive endoderm differentiation

BRD3 is dispensable for maintenance of ESC pluripotency (Di Micco et al., 2014), but its role in endoderm differentiation is not clear. We used the CRISPR/Cas system to generate homozygous hESCs lines deficient in BRD3 (*BRD3*^−/−^) (Figure 4A and S4) by inserting a puromycin (puro) resistance cassette followed by a stop codon after the first amino acid of BRD3. As a negative control, we used the CRISPR/Cas system to insert a puro resistance cassette into the *AAVS1* locus. qRT-PCR analysis showed that mRNA levels of *OCT4* and *NANOG* are comparable between *BRD3*^−/−^ hESCs and control cells (Figure 4B), suggesting that pluripotency was not affected by the loss of BRD3.

**Figure 4.**
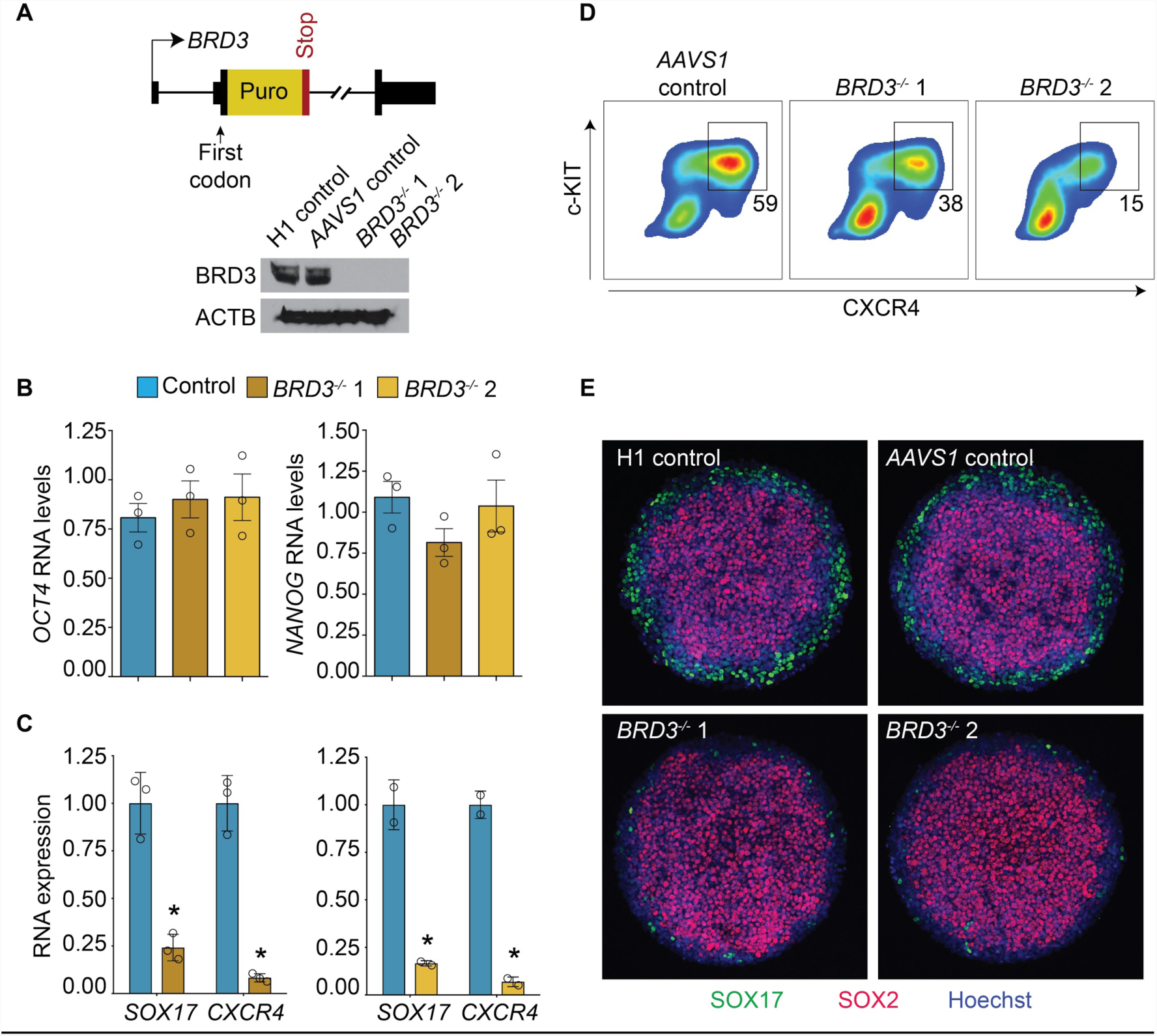
BRD3 regulates endoderm genes. **(A)** Strategy for depletion of BRD3 in hESCs using the CRISPR/Cas system and a homology plasmid to insert a puromycin resistance cassette and stop codon into the first exon of *BRD3* (top). Immunoblot was performed on cell lysates using an anti-BRD3 antibody. Samples were tested from unmodified H1 hESCs (H1 Control), hESCs in which a puromycin cassette was inserted into the *AAVS1* locus (*AAVS1* Control), and two independently-derived *BRD3*^−/−^ lines (1 and 2). ACTB was used as a loading control. **(B)** qRT-PCR was performed to quantify *OCT4* (*POU5F1*) and *NANOG* in control hESCs and two independently-derived *BRD3*^−/−^ lines. **(C)** qRT-PCR was performed to quantify *SOX17* and *CXCR4* using control hESCs and two independently-derived *BRD3*^−/−^ lines. Gene expression was analyzed after 4 days of directed differentiation. * indicates p <0.05. **(D)** Flow cytometry was performed to quantify expression of CXCR4 and c-KIT in *AAVS1* control cells and two independently-derived BRD3^−/−^ lines. The analysis was performed after 4 days of directed differentiation. Percentages of definitive endoderm cells are indicated under the boxes. **(E)** The indicated hESCs were differentiated by treatment with BMP4 for 2 days on micropatterned slides. Immunofluorescent staining was performed for SOX2 (red) and SOX17 (green) and nuclei were counterstained with Hoechst (blue).

We next differentiated *BRD3*^−/−^ hESCs toward definitive endoderm. qRT-PCR showed that mRNA levels of *SOX17* and *CXCR4*, markers of definitive endoderm, are significantly reduced with differentiation of *BRD3*^−/−^ cells compared to controls (Figure 4C). To provide further evidence that the loss of BRD3 inhibited definitive endoderm differentiation, we performed flow cytometry to quantify the production of definitive endoderm by co-expression of CXCR4 and c-KIT. We again observed a decrease in the definitive endoderm population in *BRD3*^−/−^ cells (Figure 4D). For these differentiation experiments (Figure 4C and 4D), activin treatment was used to direct the differentiation of hESCs into definitive endoderm (D’Amour et al., 2005). We then evaluated the necessity of BRD3 under other differentiation conditions. We cultured *BRD3*^−/−^ and control hESCs on micropatterned slides in the presence of BMP4 to induce differentiation of ectoderm, mesoderm, and endoderm (Deglincerti et al., 2016a; Warmflash et al., 2014). Under these conditions, differentiation is organized spatially, with ectoderm (SOX2) in the central region, definitive endoderm (SOX17) on the periphery, and mesoderm between ectoderm and endoderm. *BRD3*^−/−^ cells showed reduced SOX17 expression on the periphery of differentiating colonies as well as an expansion of central cells staining for SOX2 (Figure 4E). These results demonstrate that loss of BRD3 inhibits endoderm differentiation, a phenotype also observed with depletion of *DIGIT* (Daneshvar et al., 2016).

### BRD3 interacts with acetylated H3K18

Individual bromodomains of the BRD3 protein have been shown to interact with acetylated histones (Filippakopoulos et al., 2012), but the affinity of full-length BRD3 protein for modified histones has not been investigated. To understand how full-length BRD3 regulates genomic targets, we asked which histone modifications are recognized by BRD3. We hybridized the long and short isoforms of recombinant BRD3 protein to a histone modification peptide array and found that BRD3 shows the strongest interaction with acetylated histone H3 lysine 18 (H3K18ac) (Figure 5A, 5B, S5A, and S5B). This interaction was not detected with either bromodomain of BRD3 alone (Filippakopoulos et al., 2012).

**Figure 5.**
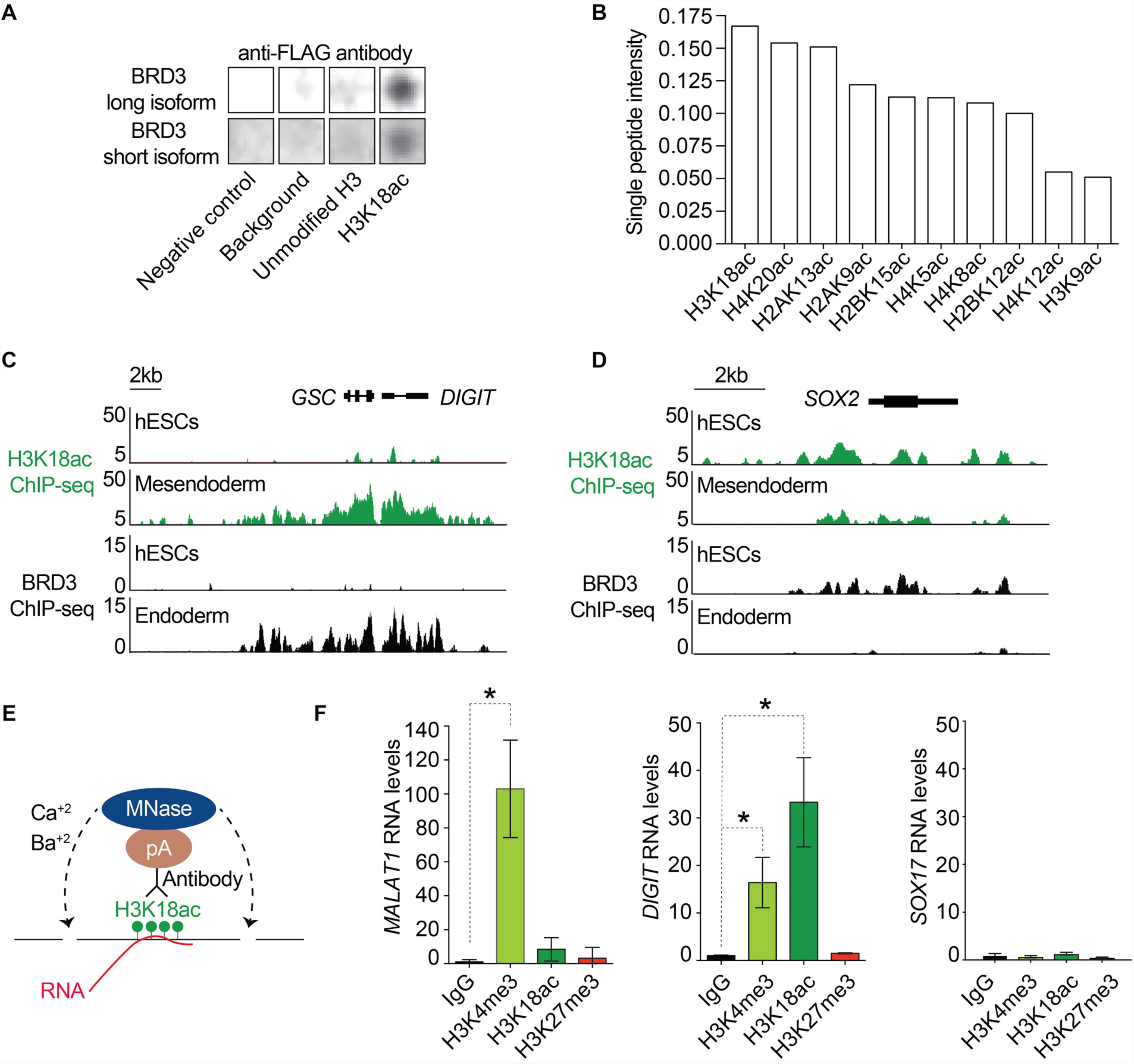
BRD3 and *DIGIT* interact with acetylated H3K18. **(A)** Binding of recombinant BRD3 to H3K18ac on a peptide modification array. Two replicates are shown, and the full arrays are shown in Figure S5A. **(B)** The top 10 modifications bound by recombinant BRD3 protein (long isoform). **(C)** and **(D)** show BRD3 occupancy (black) and H3K18ac (green) in hESCs and in mesendoderm (H3K18ac) and endoderm (BRD3) differentiation at *GSC/DIGIT* and *SOX2*, markers of definitive endoderm and pluripotency respectively. Y-axis in the ChIP-seq tracks denote normalized read count. **(E)** Schematic shows the strategy for CUT&RUNER, which takes advantage of targeted Protein A (pA) fused MNase digestion of chromatin fragments followed by isolation of the chromatin-bound RNA. **(F)** qRT-PCR shows the levels of *MALAT1* (left), *DIGIT* (middle), and *SOX17* (right) isolated by performing CUT&RUNER in hESCs differentiated toward definitive endoderm. RNA levels are normalized to *GAPDH* expression. * indicates p <0.05.

### BRD3 and H3K18ac co-occupy genes that regulate pluripotency and differentiation

Because BRD3 preferentially binds H3K18ac *in vitro*, we wanted to determine if this interaction also extends to the native chromatin in live cells. We performed ChIP-seq for BRD3 in hESCs and after 4 days of endoderm differentiation and compared occupancy of BRD3 at the *DIGIT/GSC* locus (Figure 5C) to acetylation of H3K18 (Dixon et al., 2015). We observed no evidence of BRD3 occupancy or H3K18ac at this locus in hESCs. However, BRD3 occupies this locus in endoderm. Mesendoderm gives rise to endoderm, and many endoderm genes, including *GSC*, are first activated in mesendoderm (Tada et al., 2005; Vallier et al., 2009). In mesendoderm, we observed H3K18ac in the same regions of the *DIGIT/GSC* locus that are occupied by BRD3 in endoderm. In contrast, BRD3 occupied regions of H3K18ac in hESCs at the gene encoding the pluripotency factor SOX2, and both BRD3 occupancy and H3K18ac are reduced with endoderm/mesendoderm differentiation (Figure 5D). These findings show that BRD3 and H3K18ac co-occupy genes in hESCs and with differentiation.

### DIGIT is enriched in regions of chromatin modified by H3K18ac

BRD3 localizes to actively transcribed regions of chromatin marked by H3K18ac and it physically interacts with *DIGIT*, suggesting that *DIGIT* localizes to actively transcribed regions of chromatin. We adapted the Cleavage Under Targets and Release Using Nuclease (CUT&RUN) (Skene and Henikoff, 2017) approach to isolate RNA molecules that interact with specific histone modifications (<u>C</u>leavage <u>U</u>nder <u>T</u>argets and <u>R</u>elease <u>U</u>sing <u>N</u>uclease to Assess <u>E</u>nrichment of <u>R</u>NA, CUT&RUNER) (Figure 5E). The CUT&RUN protocol uses micrococcal nuclease (MNase) to excise DNA at targeted regions. MNase function is dependent on calcium cations, and it can target both DNA and RNA molecules. However, its RNase activity can be inhibited by addition of heavy metal cations without significantly compromising its DNase activity (Cuatrecasas et al., 1967). We tested two heavy metal cations (barium and strontium), which have been reported to inhibit the RNase activity of MNase. We determined that both barium and strontium cations can inhibit the RNase activity of MNase (Figure S5C and S5D). We then used our modified approach to test enrichment of the lncRNA *MALAT1*, which interacts with chromatin at sites enriched for H3K4me3, a marker of active promoters (West et al., 2014). Results of CUT&RUNER show that *MALAT1* transcripts are enriched at regions modified by H3K4me3 compared to H3K18ac or the repressive mark H3K27me3. We then quantified enrichment of *DIGIT* in regions with H3K4me3 and H3K18ac. *DIGIT* demonstrates the greatest enrichment in regions modified by H3K18ac (Figure 5F). Enrichment was also observed in regions modified by H3K4me3, but not with IgG control or regions modified by H3K27me3. We do not find enrichment for *SOX17* mRNA in any of these regions. These results show that BRD3 and *DIGIT* are both enriched in regions of chromatin modified by H3K18ac.

### BRD3 promotes gene expression through occupancy of enhancers

BRD3 primarily occupies promoters in hESCs and shows increased occupancy of enhancers with endoderm differentiation (Figure 6A and S6A). In both cell types, BRD3 occupies genes enriched in DNA binding and transcription activity (Figure 6B). *DIGIT* is induced with endoderm differentiation and interacts with BRD3, and we investigated the overlap between genes occupied by BRD3 and those regulated by *DIGIT*. We identified 1337 genes occupied by BRD3 in endoderm differentiation and compared this to the genes inhibited in *DIGIT*-deficient cells during endoderm differentiation (Daneshvar et al., 2016). Fifty-eight genes were occupied by BRD3 in endoderm differentiation and depleted with the loss of *DIGIT*, including the key endoderm and mesendoderm genes *GSC, FOXA2*, and *EOMES* (Figure 6C and 6D). We also observed that these regulatory genes are located in larger domains occupied by BRD3, and genes in these larger domains are enriched in pathways of embryonic morphogenesis and pattern specification (Figure S6B). These results suggest that upon endoderm differentiation, BRD3 shifts to preferentially occupy enhancers, including those that regulate genes that control endoderm fate.

**Figure 6.**
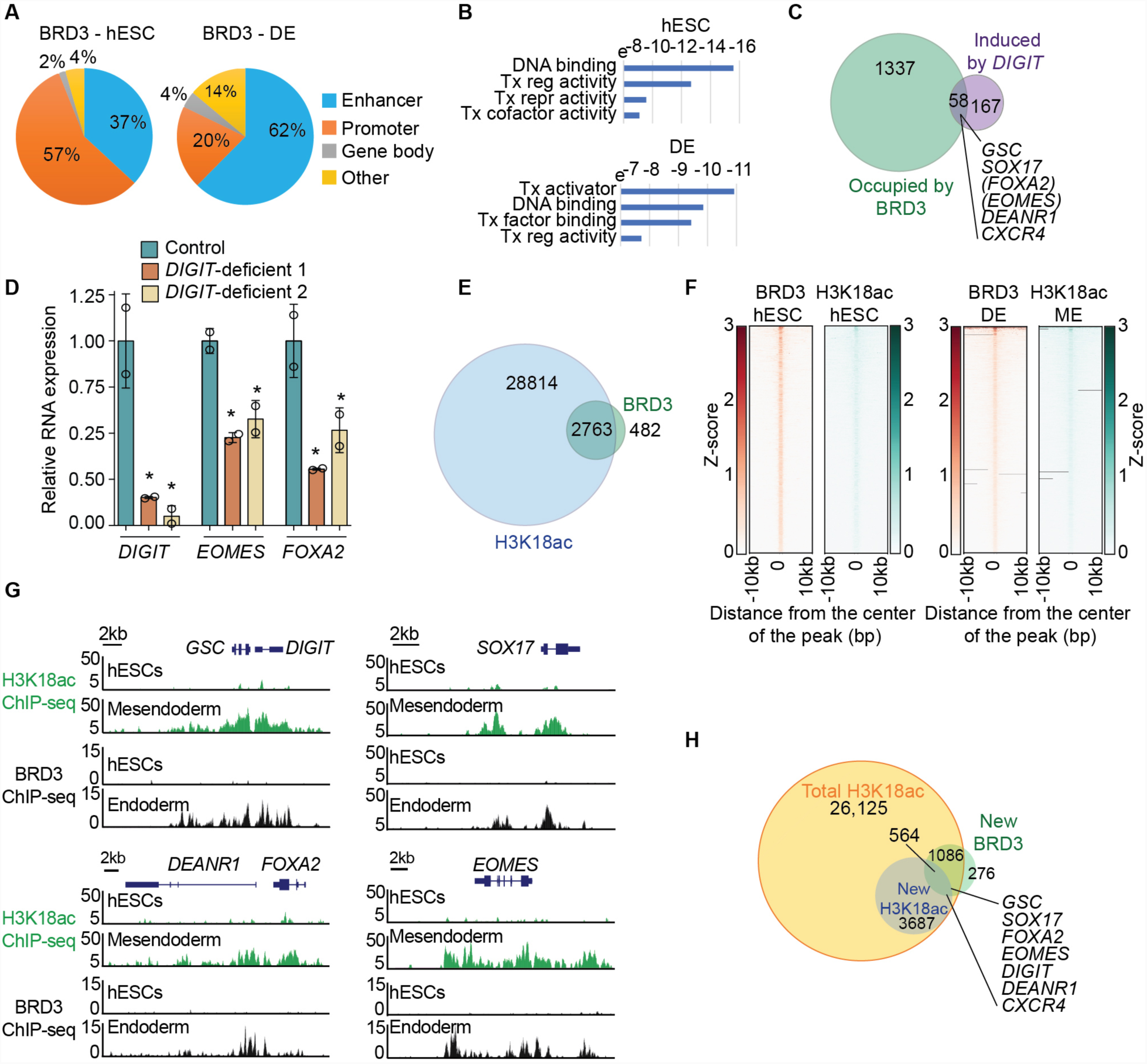
BRD3 interacts with *DIGIT*-regulated genes on H3K18ac-marked chromatin. **(A)** Genomic distribution of BRD3 in hESCs and after 4 days of definitive endoderm (DE) differentiation. **(B)** Gene ontology analysis for genes occupied by BRD3. Bars indicate p-value. Tx: transcription. **(C)** Venn diagram showing the overlap of genes occupied by BRD3 and repressed with depletion of *DIGIT* (induced by *DIGIT*) during endoderm differentiation. Endoderm genes that are regulated by *DIGIT* and occupied by BRD3 are listed. Genes in parentheses did not meet statistical significance on RNA-seq (FDR <0.01, (Daneshvar et al., 2016) and were validated by qRT-PCR in panel D. * indicates p <0.05. **(D)** qRT-PCR was performed in two independently-derived *DIGIT*-deficient hESCs lines (Daneshvar et al., 2016) after day 4 of endoderm differentiation. RNA expression is normalized to 1.0 in control cells (green) and compared to two different *DIGIT*-deficient lines. **(E)** Venn diagram showing the overlap of regions modified by H3K18ac in mesendoderm and occupied by BRD3 in endoderm differentiation. **(F)** Heatmaps show co-occupancy of BRD3 and H3K18ac in hESCs and definitive endoderm/mesendoderm (DE/ME). H3K18ac ChIP-seq analysis was performed on day 2 of mesendoderm differentiation induced by BMP4 and activin (Dixon et al., 2015). **(G)** Example mesendoderm and definitive endoderm genes that are modified by H3K18ac and occupied by BRD3 upon differentiation of hESCs toward mesendoderm and endoderm. **(H)** Venn diagram showing all regions modified by H3K18ac (yellow), newly modified by H3K18ac in mesendoderm differentiation (blue), and newly occupied by BRD3 with endoderm differentiation (green). The intersection of newly modified H3K18ac regions and newly occupied BRD3 regions contained the indicated endoderm factors.

### BRD3 and H3K18ac are associated across the genome

We previously found that BRD3 interacts with H3K18ac, and that BRD3 occupies the gene encoding SOX2 in hESCs and the gene encoding GSC with endoderm differentiation, each at sites modified by H3K18ac (Figure 5A-D). We next investigated how BRD3 and H3K18ac interact across the genome. We compared sites of H3K18ac in mesendoderm differentiation (Dixon et al., 2015) with BRD3 occupancy during endoderm differentiation, as many endoderm genes are first activated with formation of mesendoderm (Tada et al., 2005; Vallier et al., 2009). Despite analyzing occupancy under slightly different conditions of differentiation, we found that over 85% of regions occupied by BRD3 overlap with regions modified by H3K18ac (Figure 6E). In both hESCs and with endoderm differentiation, regions occupied by BRD3 are modified by H3K18ac (Figure 6F and 6G). In addition, approximately 15% of H3K18ac regions are unique in mesendoderm compared to hESCs, and 564 of these mesendoderm regions are also newly occupied by BRD3 with endoderm differentiation. This group of genes newly marked by H3K18ac and BRD3 with differentiation again includes the key endoderm genes that are also regulated by *DIGIT* (Figure 6H). These results suggest that during differentiation, BRD3 shifts to regions newly modified by H3K18ac, which encompass genes regulated by *DIGIT*.

### DIGIT recruits BRD3 to endoderm genes

The previous results suggest that *DIGIT* may act to recruit BRD3 to sites of H3K18ac to induce expression of endoderm genes. To test this hypothesis, we differentiated *DIGIT*-deficient (*DIGIT*^*gfp/gfp*^) and wildtype hESCs toward endoderm to determine how BRD3 occupancy was affected by depletion of *DIGIT*. We found that BRD3 occupancy at *GSC* and *FOXA2* (Figure 6G) was impaired with depletion of *DIGIT* (Figure 7A). As a control, we also assessed BRD3 occupancy at *HEXIM2*, a gene occupied by BRD3 in hESCs and with endoderm differentiation but not affected by depletion of *DIGIT* (Figure S7). In contrast to *GSC*, where we observed reduced BRD3 occupancy in *DIGIT*-deficient cells, BRD3 occupancy at *HEXIM2* was not reduced (Figure 7B). These results suggest that *DIGIT* is required for recruitment of BRD3 to endoderm genes (Figure 7C).

**Figure 7.**
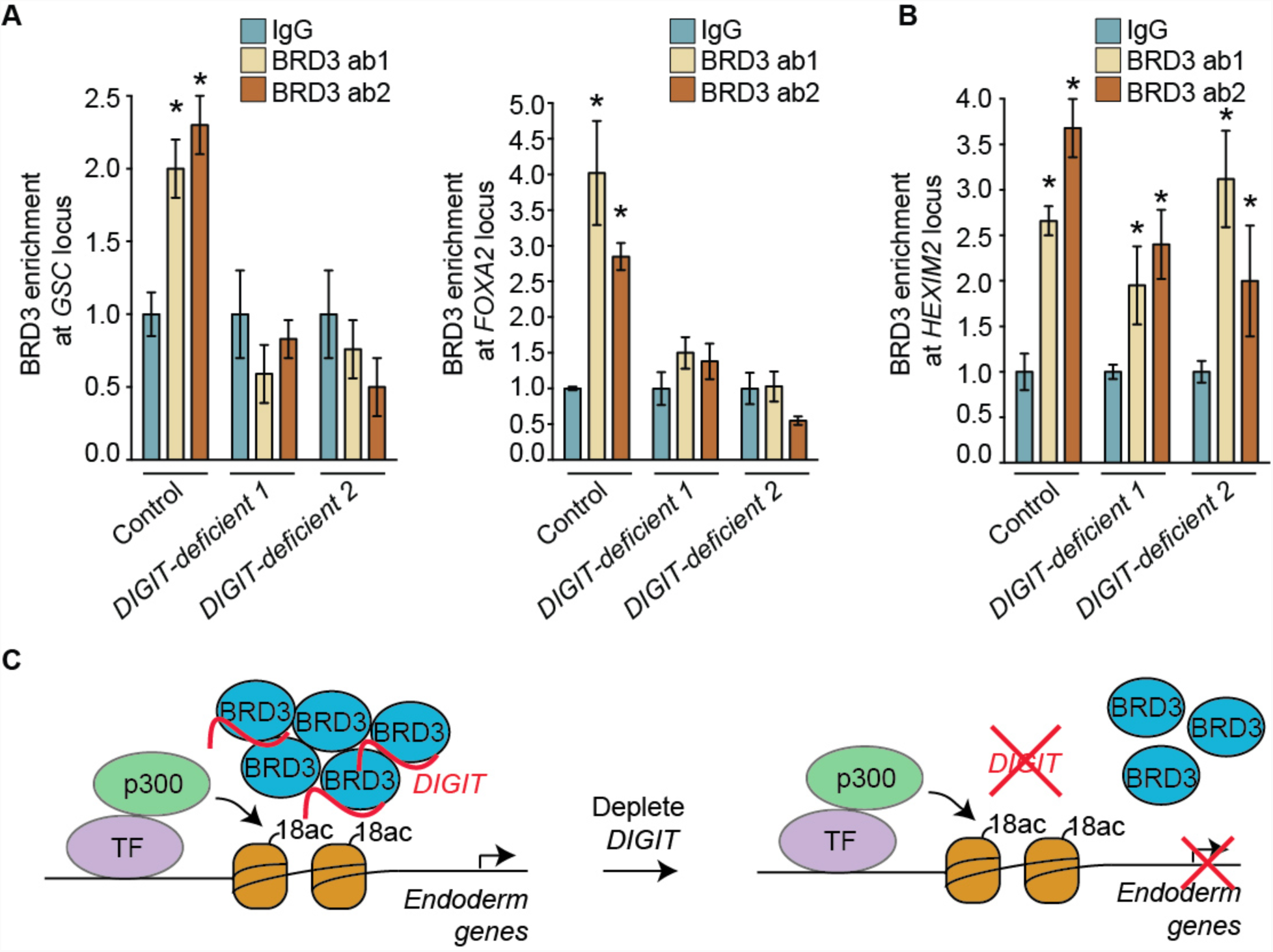
Loss of *DIGIT* inhibits recruitment of BRD3. **(A)** CUT&RUN was performed followed by qPCR to quantify enrichment of BRD3at the *GSC* locus (left) and *FOXA2* locus (right). Chromatin release was directed using two antibodies recognizing BRD3 (ab1 and ab2). Two independently-derived *DIGIT*-deficient lines (*DIGIT*^*gfp/gfp*^) were used for each experiment. **(B)** CUT&RUN was performed as in (A) using primers that recognize *HEXIM2*. * indicates p <0.05, as calculated by ANOVA and post hoc test, compared to IgG. **(C)** Schematic showing condensates of BRD3 and *DIGIT* interacting with H3K18ac at the enhancer of an endoderm gene to induce gene expression. H3K18 is acetylated by CBP/p300 following recruitment by transcription factors (TF) (Ito et al., 2000; Jin et al., 2011). In the absence of *DIGIT*, BRD3 is not able to bind the enhancer, and the endoderm gene is not activated.

## Discussion

Many lncRNAs interact with proteins to regulate gene expression. The interactions initially identified were found to repress gene expression, in the case of *XIST* through the interaction with YY1 (Jeon and Lee, 2011) and for *HOTAIR* and *ANRIL* through interaction with the polycomb repressive complexes (PRC) (Rinn et al., 2007; Yap et al., 2010). Additional investigations revealed that lncRNAs also partner with proteins to induce gene expression (Bose et al., 2017; Postepska-Igielska et al., 2015). Here we demonstrate that *DIGIT* interacts with BRD3 proteins in phase-separated condensates to regulate gene expression and endoderm differentiation. BRD3 binds H3K18ac, and during endoderm differentiation, BRD3 shifts to occupy enhancers within domains of H3K18ac. BRD3 can co-occupy genes with H3K18ac independent of *DIGIT* (Figure 7B and S7), but BRD3 requires *DIGIT* to occupy regions of H3K18ac near a specific subset of genes (Figure 7A and 7C). Thus, the interaction with *DIGIT* determines the specificity of a set of BRD3 targets, through which BRD3 regulates endoderm differentiation.

*DIGIT* and BRD3 cooperate to promote expression of key endoderm genes. *DIGIT* is divergently transcribed from *GSC*, and activation of *GSC* can, at least partially, rescue endoderm differentiation (Daneshvar et al., 2016). Results from this study show that *DIGIT* interacts with BRD3 to control *GSC* expression and suggest that the interaction between *DIGIT* and BRD3 also regulates additional endoderm and mesendoderm genes. Thus, through interaction with BRD3, *DIGIT* expands its influence beyond the immediate vicinity where it is transcribed.

BRD4 forms phase-separated condensates in association with Mediator to regulate gene expression (Sabari et al., 2018), and BRD2, BRD3, and BRD4 each form distinct phase-separated condensates during endoderm differentiation. *DIGIT* preferentially interacts with BRD3, but there are likely to be other lncRNA partners of BRD3, as well as lncRNAs that preferentially interact with BRD2 and BRD4. The effect of lncRNA-BET family protein interactions on the formation or stability of phase-separated condensates is unclear and will need to be addressed in future experiments. The lncRNA *NEAT1* interacts with the protein NONO to promote phase-separated paraspeckles (Yamazaki et al., 2018), and the lncRNA *MALAT1* is found in phase-separated nuclear speckles (Hutchinson et al., 2007). The interaction between *DIGIT* and BRD3 now demonstrates a role for lncRNAs in transcriptional phase-separated condensates.

BRD4 preferentially occupies enhancers in hESCs, while BRD2 and BRD3 preferentially occupy promoters (Di Micco et al., 2014; Engelen et al., 2015). We observe this same bias for BRD3 occupancy at promoters in hESCs, where BRD3 is dispensable. However, with endoderm differentiation, where BRD3 is required, BRD3 shifts to enhancers. These findings suggest enhancer occupancy may be a path through which BRD3 activates developmental regulators and BET family proteins regulate gene expression.

BRD3 interacts most intensely with H3K18ac, a modification catalyzed by CBP/p300 (Jin et al., 2011). CBP/p300 is also responsible for H3K27ac, but H3K27ac showed <10% enrichment for BRD3 binding compared to H3K18ac (data not shown), suggesting that BRD3 has minimal affinity for H3K27ac. In mouse endoderm differentiation, activin signaling leads to increased H3K18ac at the *Gsc* gene (Xi et al., 2011). Acetylation of H3K18 at *Gsc* requires both Smad4 and TRIM33, each of which interacts with Smad2/3 in response to activin signaling. These findings suggest that genome occupancy by Smad2/3 at new enhancers leads to recruitment of CBP/p300 (Janknecht et al., 1998; Pouponnot et al., 1998), induction of H3K18ac, and binding of of the BRD3-*DIGIT* complex to drive expression of *Gsc* and other Smad2/3 target genes.

BRD2, BRD3, and BRD4 are ubiquitously expressed and have non-overlapping functions in many cell types (Houzelstein et al., 2002; Paillisson et al., 2007; Shang et al., 2009). Our findings indicate that the interaction between BET family proteins and lncRNAs help specify gene targets. These findings suggest that interactions with different lncRNAs may be responsible for determining different subsets of genes regulated by BET family proteins in different cell types. While we propose that BET family proteins may bind several different lncRNAs species, this binding is limited, as we do not see evidence of broad interaction between BRD3 and polyA RNAs. The interaction with BET family proteins is also not limited to polyA RNAs, as enhancer (e) RNAs can also bind BET family proteins (Rahnamoun et al., 2018). It remains unclear how interactions between BET family proteins and noncoding RNAs regulate gene expression. BRD4 mediates pause release (Liu et al., 2014), while BRD2 and BRD3 show nucleosome remodeling activity to promote processivity of RNA polymerase (LeRoy et al., 2008). Depletion of *DIGIT* suggests that one mechanism of regulation may be at the level of recruitment of BET family proteins, but interaction with lncRNAs could also regulate the activity of BET family proteins. Further studies are needed to understand the full breadth of interactions between BET family proteins and lncRNAs and how these interactions regulate gene expression, mechanisms which likely control numerous cell functions in development and disease.

## Materials and Methods

### Embryonic stem cell culture and differentiation

H1 (WA01) cells (WiCell) were cultured in feeder-free conditions in mTeSR1 (Stem Cell Technologies) and passaged by Accutase (Stem Cell Technologies) as described (Daneshvar et al., 2016). Endoderm differentiation was induced with 50 ng/mL Activin A (R&D Systems) in RPMI with B27 supplement (Thermo Fisher Scientific). Differentiation of hESCs on micro-patterned slides was performed as described (Deglincerti et al., 2016b; Warmflash et al., 2014). Briefly, hESCs were dissociated into single cells using Accutase and plated on micropatterned slides (Cytoo) that were coated with laminin-521 (BioLamina) as described. Differentiation toward germ layers was induced by addition of 50 ng/mL of BMP4 (R&D Systems) to mTeSR1 media. Cells were fixed and immunostained. Immunofluorescence microscopy was performed on an EVOS FL Cell Imaging System (Thermo Fisher Scientific).

### S1m aptamer-assisted RNA pull-down

Plasmids expressing *DIGIT, DIGIT-4xS1m*, and scramble-*DIGIT* (*SCRM-4xS1m*) were transiently transfected into 3x10^6^ H1 cells by lipofectamine 3000 (Thermo Fisher). Cells were differentiated toward definitive endoderm for three days. Cells were washed with DPBS and irradiated with 400 mJ of energy on ice in a UV cross-linker (Stratalinker). Nuclei of the UV cross-linked cell were isolated by suspending the cells in a hypotonic buffer (20 mM Tris-HCl pH 7.4, 10 mM NaCl, 3 mM MgCl_2_) for 15 minutes, adding NP-40 to a final concentration of 0.5%, and then centrifuging for 10 minutes at 850 *g*. Nuclei were lysed in lysis buffer (150 mM KCl, 25 mM Tris-HCl pH 7.4, 5 mM EDTA, 5 mM MgCl_2_, 1% NP-40, 0.5 mM DTT, Roche mini-tablet protease inhibitor, and 100 U/mL RNAseOUT), and debris was cleared by centrifugation at 16000 *g*. The supernatant containing the nuclear lysate was then cleared with 50 μL of Avidin Agarose beads (Thermo Fisher, catalog number: 20219) to deplete the lysate from biotin (Leppek and Stoecklin, 2014). The cleared lysate was incubated with 150 μL of Streptavidin C Dynabeads (Thermo Fisher) for 4 hours on a rocker at 4°C. Beads were collected using a magnet and washed three times with a wash buffer of the same composition as the lysis buffer, except KCl was increased to 350 mM. To release the proteins, beads were incubated with RNase A and resuspended in 2x LDS sample buffer. The entire lysates were resolved on a polyacrylamide gel. Segments of the gel between 30 to 200 kDa were excised and analyzed by mass spectrometry. Mass spectrometry was performed at Harvard Medical School Taplin Mass Spectrometry Core. Analysis of the proteomic data was performed using the Crapome proteomic analysis pipeline (Mellacheruvu et al., 2013).

### Pull-down of cross-linked polyadenylated RNA and protein complexes

The pull-down of polyadenylated RNAs was performed as described before (Castello et al., 2016). Ten million hESCs were differentiated toward endoderm for 3 days. Cells were UV cross-linked as described above. The nuclear fraction was isolated as described above. Nuclei were lysed in lysis/binding buffer (20 mM Tris-HCl pH 7.5, 500 mM LiCl, 0.5% LiDS, 1 mM EDTA, 5 mM DTT). The lysate was then incubated with 200 μL of oligo-dT 25 Magnetic Beads (NEB, S1419S) for 10 minutes. The beads were washed two times with wash buffer 1 (20 mM Tris-HCl pH 7.5, 500 mM LiCl, 0.1% LiDS, 1 mM EDTA, 5 mM DTT), two times with wash buffer 2 (20 mM Tris-HCl pH 7.5, 500 mM LiCl, 1 mM EDTA), and one time with low salt buffer (20 mM Tris-HCl pH 7.5, 200 mM LiCl, 1 mM EDTA). Proteins were released by addition of 5 μg of RNase A (Sigma) in RNA digestion buffer (20 mM Tris-HCl pH 7.5, 30 mM NaCl, 5 mM MgCl2, 2 mM DTT, 1 tablet/10 mL Mini Complete Protease Inhibitors, EDTA-free). For RNase digested control, RNase A was added to the low salt buffer wash.

### Transfection and genome editing

For transfection, hESCs were dissociated by Accutase and resuspended in buffer 1M nucleofection buffer (5 mM KCl, 15 mM MgCl_2_, 120 mM Na_2_HPO_4_/NaH_2_PO_4_ pH 7.2, 50 mM Mannitol) and transfected in a Nucleofector 4D (Lonza) (Chicaybam et al., 2013). For genome editing, 3 μg of the homology construct and 1 μg of the Cas9/gRNA construct were co-transfected. For the generation of BRD3 knockout lines, cells were treated with 0.5 μg/mL puromycin for 8 days starting 48 hours after transfection. Puromycin-resistant colonies were picked and transferred to 48-well plates. For insertion of mEGFP into BRD3, 72 hours after transfection of constructs, cells were sorted by flow cytometry using a FACSAria flow cytometer. For the generation of hESCs that stably express NLS-mEGFP, we inserted a DNA sequence expressing NLS-mEGFP under an EF1α promoter and puromycin resistance gene under an RPBSA into the H1 genome using the Sleeping Beauty Transposon System (Kowarz et al., 2015). Cells were treated with puromycin for 8 days and expression of NLS-mEGFP was confirmed by fluorescence microscopy.

### Expression and purification of FLAG-BRD3, FLAG-mEGFP-BRD3, and FLAG-mEGFP proteins

The cDNA for the short isoform of human BRD3 (Dharmacon, clone ID: 4856840) was used to generate the long isoform of BRD3. The extra sequence was generated by PCR amplification from hESC genomic DNA and added by Gibson assembly (NEB). The long isoform of BRD3 with N-terminal mEGFP (FLAG-mEGF-BRD3) and the short isoform of BRD3 (FLAG-BRD3) were subcloned into pFastBac1 Baculovirus expression plasmid, each with a single N-terminal FLAG-tag. The FLAG-mEGFP-BRD3 fusion protein containing the long isoform of BRD3 was created with a 7 amino acid linker sequence (FLAG-GSAAAGS-mEGFP-GSAAAGS-BRD3). The FLAG-BRD3 fusion created using the short isoform of BRD3 was created without a linker between the FLAG sequence and the start of BRD3.

One liter of Sf9 cells infected with P3 viral particles expressing each construct was grown in ESF 921 medium (Expression Systems) at 28°C. Forty two hours post infection, FLAG-tagged proteins were affinity purified from the nuclear extract as described (Abmayr et al., 2006; Grau et al., 2011). All steps were carried out at 4°C. In brief, 500 μL of equilibrated anti-FLAG M2 affinity gel (50% slurry, Sigma) was added to isolated nuclei in BC600 (20 mM HEPES pH 7.9, 20% glycerol, 1.5 mM MgCl_2_, 0.6 M KCl, 0.2 mM EDTA, 0.2 mM PMSF, 0.5 mM DTT, 0.05% NP-40, and Complete Protease Inhibitor Cocktail (Roche)), after 3-4 hrs the beads were passed through Econo-Pac chromatography columns (Bio-Rad) and washed with 20x bead volume of wash buffer in the following order: BC600(x2), BC1200, BC600, BC300 and eluted twice by incubation with 250 μL of 400 μg/mL FLAG peptide in BC300 (without NP-40). Each elution was carried out for 20 min. Eluted protein was concentrated using Amicon Ultra centrifuge filters. The concentration of the complex was determined by Bicinchoninic acid (BCA) assay (Thermo Fisher).

FLAG-mEGFP (FLAG-GSAAAGS-mEGFP-GSAAAGS) was also cloned into pFastBac1 and purified as follows. After 42 hrs of infection with P3 viral stock, cells were lysed and flash-frozen in BC500 (50 mM Tris pH 8.0, 0.5 M NaCl, 20% glycerol, 0.05% NP-40, 0.2 mM EDTA, 0.2 mM PMSF, 0.5 mM DTT, and Roche complete protease inhibitor tablets). The frozen extract was thawed, spun down at 14,000 *g* for 30 min, and the supernatant was incubated with 500 μL of equilibrated M2-agarose beads for 3-4 hrs. Purification was performed using Econo-Pac chromatography columns as described above and elsewhere (Grau et al., 2011).

### Fluorescence recovery after photobleaching (FRAP)

For FRAP experiments, hESCs were grown and differentiated on coverslip bottom tissue culture dishes (FluoroDish, World Precision Instruments) and imaged using a Nikon A1R confocal inverted microscope. Photobleaching was performed using the 488nm laser at 95% power. A circular region of interest (ROI) with a radius of 1.5 μm was defined around a single BRD3 punctum. Images were acquired at indicated times. The fluorescence intensity of the puncta were measured by ImgeJ. As a control experiment, the same setup was used except that the photobleaching step was eliminated.

### Histone modification arrays

Modified Histone arrays (Active Motif) were used as described (Saltzman et al., 2018) to identify the histone targets of BRD3. The arrays were washed one time in PBST (PBS, 0.05% v/v Tween-20), blocked with skim milk (5% w/v in PBST) for 1 hour, and washed with BSA (3% w/v in PBST). Arrays were incubated overnight at 4°C in binding buffer (PBST, 0.45% w/v BSA, 0.5 mM EDTA, 0.1 mM DTT, 10% v/v glycerol) with 100 nM of each protein (FLAG-mEGF-BRD3 long isoform, FLAG-BRD3 short isoform, or FLAG-mEGFP). Arrays were then probed with the primary (anti-FLAG) and secondary (anti-mouse IgG) antibodies. The arrays were incubated in Western Blotting Luminol Reagent (Santa Cruz) and exposed to autoradiography film. Developed films were scanned and analyzed with ArrayAnalyze software (Active Motif).

### Chromatin immunoprecipitation (ChIP)

Chromatin immunoprecipitation was performed as described (Daneshvar et al., 2016). Approximately 6 million cells per IP were cross-linked with formaldehyde and sheared using a Covaris sonicator. IPs were performed with 1.5 μg of antibody per IP.

### Cleavage Under Targets and Release Using Nuclease (CUT&RUN) to quantify DNA enrichment

CUT&RUN was performed as described (Skene and Henikoff, 2017; Skene et al., 2018) with following modifications: 1) Dissociated cells were cross-linked by resuspension in a cross-linking buffer (11% Formaldehyde, 0.1 M NaCl, 1 mM EDTA, 0.5 mM EGTA, 50 mM HEPES) for 10 minutes and quenched in 125 mM glycine in DPBS. 2) Tween-20 was added to all wash buffer at a final concentration of 0.2%. 3) Bovine Serum Albumin (BSA) was added to wash buffers at the final concentration of 0.25% w/v. 4) CUT&RUN fragments were reverse cross-linked in 65°C for 6 hours before DNA extraction. Recombinant Protein A fused to MNase was a gift from Steven Henikoff (Fred Hutchinson Cancer Research Center).

### Cleavage Under Targets and Release Using Nuclease to isolate chromatin-associated RNA (CUT&RUNER)

This assay was performed similarly to cross-linked CUT&RUN, as described above, with following modifications: 1) Superase-In RNase inhibitor (Thermo Fisher) was added to all buffers at a final concentration of 20 U/mL. 2) In the targeted digestion step, barium chloride was added at a final concentration of 10 mM along with CaCl_2_ at a final concentration of 2 mM. 3) RNase A was removed from the 2x STOP buffer. 4) SDS was removed from the reverse cross-linking step to avoid its precipitation with barium (Putnam and Neurath, 1944).

### Immunostaining

Cells were fixed in paraformaldehyde (4% in DPBS). Cells were blocked and permeabilized in a blocking buffer (0.1% BSA, 3% normal donkey serum, 0.1% Triton X-100 in PBS). After incubation with antibodies, cells were washed three times in a wash buffer (0.1% Tween-20 in DPBS) and one time in DPBS. Nuclear staining was performed by addition of Hoechst to the last wash buffer.

### In vitro droplet assay

The concentration of the recombinant mEGFP-BRD3 and mEGFP were adjusted to varying concentrations with indicated final salt concentrations and 10% PEG-8000 (crowding agent) in Buffer D (50 mM Tris-HCl pH 7.5, 10% glycerol, 1mM DTT). The protein solution was then loaded onto a homemade chamber slide (a microscope glass slide with a coverslip attached by two parallel strips of double-sided tape). Slides were then imaged with an Andor confocal microscope with a 150x objective.

### In vitro transcription and fluorescent labeling of RNA

The cDNA encoding full-length spliced *DIGIT* was *in vitro* transcribed using the HighScribe T7 High Yield RNA synthesis kit (NEB, E2040s). Cyanine 3-UTP (Enzo Life Sciences, ENZ-42505) was added to the reaction at the final concentration of 2.5 mM. Synthesized RNA was purified and tested for integrity and fluorescence on a bleach agarose gel (Aranda et al., 2012).

### Sequential immunostaining and single molecule RNA-FISH

Cells were grown and differentiated on matrigel coated coverslips (Daneshvar et al., 2016). Cells were fixed in 3.7% formaldehyde in DPBS, followed by permeabilization in 1% Triton X-100. After staining with primary and secondary antibodies, cells were fixed in the fixation buffer one more time. Cells were then washed with smFISH wash buffer (10% v/v formamide, 2X SSC in DPBS) and hybridized with smFISH probes (Biosearch Technologies) in hybridization buffer (0.1 g/mL dextran sulfate, 10% v/v deionized formamide, 2x nuclease-free SSC, in nuclease-free water) overnight, and washed three times in wash buffer and one time in 2x SSC, before cells were stained with Hoechst and mounted on slides.

Slides were imaged on a StellarVision inverted microscope (Optical Biosystems, model SV20HT) using a Nikon CFI S Plan Fluor ELWD 0.45NA 20xC air objective. Synthetic Aperture Optics (SAO) illumination was achieved with dedicated 473 nm, 532 nm, and 660 nm laser lines (Laser Quantum) and Hoechst was excited with a 365 nm LED. Exposure times were determined for a randomly chosen area of cells near the center of the coverslip, then nine single-Z-plane fields of view (FOV) (333 μm x 333 μm) were automatically captured in an outwards clockwise spiral from the first FOV stage position. Each channel was acquired sequentially in the following order: 365 nm, 473 nm, 532 nm, 660 nm. Image-based autofocus on the DAPI channel was used to adjust focus after stage movements. Images were captured with an Andor sCMOS Zyla 4.2 Plus camera and raw image data was reconstructed using SV Recon software (Optical Biosystems, version 2.9.118). The pixel size in the final 4096 x 4096 pixel reconstructed image is 81 nm.

A total of 2206 nuclei from 1 mm^2^ area were included for analysis of *DIGIT lncRNA, GSC* mRNA, and BRD3 protein as follows. First, the centroid pixel and masked area of a total of 379 *DIGIT* and 2412 *GSC* smFISH spots were identified by SV Quant (Optical Biosystems, version 7.03) using intensity, modulation, and shape descriptor thresholds. All subsequent analysis was done in FIJI. To isolate nuclear BRD3 condensate areas for analysis, thresholded DAPI regions of interest (ROIs) were overlaid on BRD3 images and all extranuclear BRD3 signal was removed. DAPI-restricted BRD3 areas were thresholded and overlaid separately to each of the *DIGIT* or *GSC* spot centroid maps from SV Quant to determine colocalization. Colocalization was defined as having the centroid pixel of either a *DIGIT* or *GSC* smFISH spot occur inside of a BRD3 ROI.

### Plasmids and molecular cloning

Homology constructs were generated by amplification of homology arms from hESC genomic DNA and cloned into a BamHI/SalI digested PUC19 backbone by Gibson assembly (repliQ HiFi Assembly Mix, Quanta Bio). For the generation of mEGFP expressing hESCs, we first added an NLS to the mEGFP-N1 plasmid. The NLS-mEGFP cassette was then sub-cloned into the Sleeping Beauty pSBbi-Pur plasmid. mEGFP-N1 plasmid was a gift from Michael Davidson (Addgene plasmid # 54767). pX330-U6-Chimeric_BB-CBh-hSpCas9 was a gift from Feng Zhang (Addgene plasmid # 42230). pSBbi-Pur plasmid was a gift from Eric Kowarz (Addgene plasmid # 60523). pCMV(CAT)T7-SB100 was a gift from Zsuzsanna Izsvak (Addgene plasmid # 34879). The 4xS1m construct was a gift from Georg Stoecklin (Heidelberg University).

### Co-immunoprecipitation (co-IP)

Nuclei from 5 million cells were prepared as described above. Nuclei were lysed in a IP lysis/wash buffer (50 mM Tris-HCL PH 7.5, 150 mM NaCL, 1 mM EDTA, 1% Triton X-100, 1x Halt protease inhibitor). Nuclear debris was pelleted by centrifugation at 16000 *g* for 20 minutes. The supernatant was incubated with 4 μg of antibody on a rotator at 4°C for 16 hours. 40 μL of Dynabeads Protein G (Thermo Fisher, 10003D) was washed in the lysis/wash buffer, incubated in the lysate and antibody mix, and incubated for 1 hour at 4°C. Beads were precipitated by a magnetic rack and washed three times in lysis/wash buffer. Proteins were eluted by adding 2x sample buffer to the beads and boiling.

### Quantification of MNase’s DNase and RNase activity

The DNase and RNase activity of MNase in the presence of different cations was tested as described (Cuatrecasas et al., 1967). Thirty micrograms of salmon sperm DNA (Thermo Fisher, 15632011) or yeast tRNA (Thermo Fisher, AM7119) was used for MNase activity. pA-MNase was added to each reaction to the final concentration of 700 ng/mL. Reactions were incubated at 37°C. 260 nm absorbance was measured every 3 minutes by a Nanodrop 2000 Spectrophotometer (Thermo Fisher).

### RNA-immunoprecipitation (RIP)

RIP was performed as described (Rinn et al., 2007). Nuclear fraction was prepared as described above. Nuclei were then lysed in RIP buffer (150 mM KCl, 25 mM Tris pH 7.4, 5 mM EDTA, 0.5 mM DTT, 0.5% NP-40, RNase-inhibitor, and protease inhibitor). Nuclear membrane and debris were pelleted by centrifugation at 16,000 *g* for 10 minutes. 5 μg of each antibody was added to each lysate and incubated in rotator at 4°C for 3 hours. 40 μL of protein G Dynabeads (Thermo Fisher) was added to each reaction and incubated for one hour on a rotator at 4°C. Beads were precipitated on a magnetic rack and washed three times with RIP buffer. Beads were resuspended and boiled in 2x sample buffer and a reducing agent.

### Antibodies

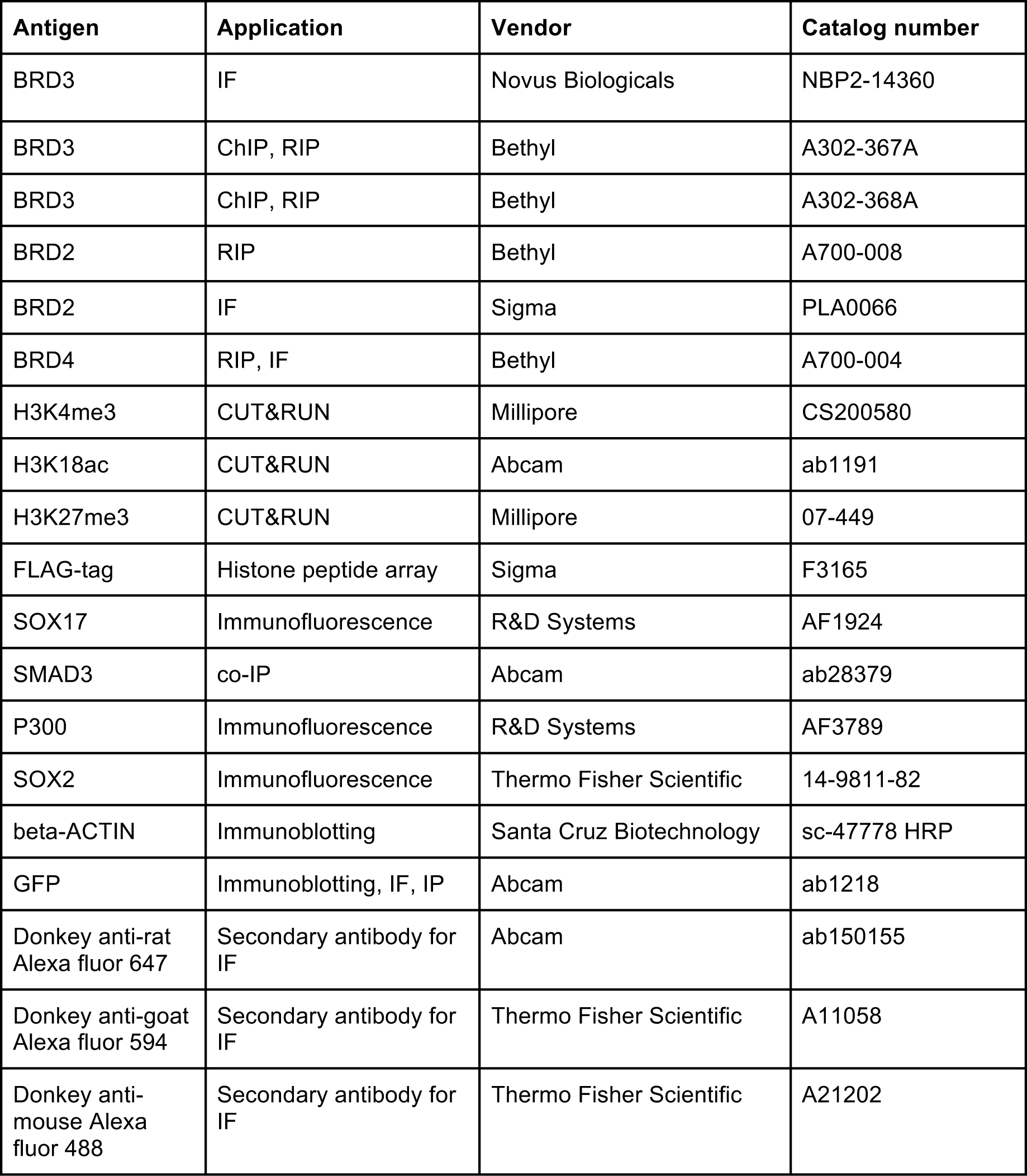

| Antigen | Application | Vendor | Catalog number |
| --- | --- | --- | --- |
| BRD3 | IF | Novus Biologicals | NBP2-14360 |
| BRD3 | ChIP, RIP | Bethyl | A302-367A |
| BRD3 | ChIP, RIP | Bethyl | A302-368A |
| BRD2 | RIP | Bethyl | A700-008 |
| BRD2 | IF | Sigma | PLA0066 |
| BRD4 | RIP, IF | Bethyl | A700-004 |
| H3K4me3 | CUT&RUN | Millipore | CS200580 |
| H3K18ac | CUT&RUN | Abcam | ab1191 |
| H3K27me3 | CUT&RUN | Millipore | 07-449 |
| FLAG-tag | Histone peptide array | Sigma | F3165 |
| SOX17 | Immunofluorescence | R&D Systems | AF1924 |
| SMAD3 | co-IP | Abcam | ab28379 |
| P300 | Immunofluorescence | R&D Systems | AF3789 |
| SOX2 | Immunofluorescence | Thermo Fisher Scientific | 14-9811-82 |
| beta-ACTIN | Immunoblotting | Santa Cruz Biotechnology | sc-47778 HRP |
| GFP | Immunoblotting, IF, IP | Abcam | ab1218 |
| Donkey anti-rat<br>Alexa fluor 647 | Secondary antibody for<br>IF | Abcam | ab150155 |
| Donkey anti-goat<br>Alexa fluor 594 | Secondary antibody for<br>IF | Thermo Fisher Scientific | A11058 |
| Donkey anti-mouse<br>Alexa fluor 488 | Secondary antibody for<br>IF | Thermo Fisher Scientific | A21202 |

### Chromatin Immunoprecipitation and sequencing (ChIP-seq) analysis

ChIP-seq was performed as previously described (Daneshvar et al., 2016) for BRD3 in hESCs and after 4 days of endoderm differentiation. ChIP-seq libraries prepared from whole cell extracts were used for background. Duplicate samples were analyzed for each condition. Two replicates of H3K18ac ChIP-seq data for hESCs (H1) and mesendoderm (ME) and control datasets were downloaded from GEO (GSE16256) (Dixon et al., 2015). We mapped the raw reads of combined replicates to human reference genome (hg19) using Bowtie2 (version 2.1.0) (Langmead and Salzberg, 2012) with the following settings: *bowtie2 -k2 -N1 -L32 --end-to-end -p 4 --phred33 -x bowtie2_index -U sample.fastq 1>sample.sam 2>sample.log*. SAM files were converted into bam format using Samtools (Li et al., 2009): *samtools view -bS sample.sam > sample.bam*. Occupied regions were called using MACS2 (Zhang et al., 2008) with the following settings: *macs2 callpeak -t DE68_2reps.bam -c DE_input.bam -f BAM -g 2.9e9 -n DE_BRD3 -q 0.01 --nomodel --shiftsize=150 -B --broad --broad-cutoff=0.1 --SPMR.* We defined regions enriched in endoderm differentiation by using Bowtie2 to call BRD3 peaks in endoderm using BRD3 occupancy in hESCs as background. We then removed any regions that were not called as occupied by BRD3 in endoderm when whole cell extract was used as a background. We performed the same analysis for H3K18ac using the data for hESCs and mesendoderm. Regions occupied by BRD3 in hESCs were defined by comparing BRD3 ChIP-seq data in hESCs to whole cell extract background. We used H3K27ac histone modification (GSE16256) to define active enhancers (Creyghton et al., 2010; Rada-Iglesias et al., 2011). H3K27ac peaks were called using the same methods as for H3K18ac ChIP-seq data. Occupied regions were assigned to a gene if the region was within 10 kilobases upstream of the TSS or overlapped the gene body.

### Visualization of regions occupied by BRD3 and modified by H3K18ac and H3K27ac

We generated peak tracks using MACS2 with the following settings: *macs2 bdgcmp -t treatment_pileup.bdg -c control_lambda.bdg -o track_logLR.bdg -m logLR -p 0.00001* and then converted the output BedGraph formated file into BigWig format based on UCSC toolkit (*bedGraphToBigWig*) (Kent et al., 2010).

### Classification of regions occupied by BRD3

BRD3 binding regions were classified into four categories based on overlaps with the following regions in the genome: (1) promoters, (2) enhancers, (3) gene body, and (4) others. Promoter regions were defined as the region from upstream 2kb to downstream 2kb of transcription start sites (TSS). Enhancers were defined by the presence of H3K27ac that did not overlap promoter regions. Gene body was defined from the TSS to transcription termination for each gene annotated in UCSC. Regions were first assigned to promoters. If a region did not overlap by at least 1 bp with a promoter region, the region was then evaluated for overlap with enhancer regions. If a region did not overlap a promoter or an enhancer, it was evaluated for overlapping a gene body. Regions that did not overlap with any previous category were classified as other.

### Heatmaps of BRD3 and H3K18ac peaks

Heatmaps of BRD3 and H3K18ac at hESCs and endoderm/mesendoderm represent the read densities of BRD3 and H3K18ac ChIP-seq data at the regions surrounding the summits of BRD3 peaks (from upstream 10 kb to downstream 10 kb of the summits) for hESCs and endoderm/mesendoderm respectively. We converted all mapped reads in Bam format into BigWig format using the *bamCoverage* command from the *Deeptool* toolkit (version: 3.1) (Ramírez et al., 2016). We then calculated the read densities in each bin using the *computeMatrix* command from the *Deeptool* toolkit with the BigWig file as the input. We plotted the heatmaps using the *plotHeatmap* command from the *Deeptool* toolkit with the read densities output by *computeMatrix*.

### Distributions of read counts for regions occupied by BRD3 in endoderm

We used *HTseq* (v0.6.1) with the setting: (*“-r name -a 10 -i gene_id -m union <alignment_file> <gff_file>”)* (Anders et al., 2015) calculate the number of reads mapped to the regions occupied by BRD3 based on BRD3 ChIP-seq and control ChIP-seq. The read counts were normalized by subtracting the number of mapped reads of the control ChIP from the BRD3 ChIP. The distribution graph was plotted based on the lengths of BRD3 binding regions from the smallest number to the largest number.

### Gene Ontology (GO) analyses

GO functional enrichment analysis (Huang et al., 2009a, 2009b) (version 6.7) was performed using the indicated groups of BRD3-associated genes. For analysis in hESCs, the background was defined as all expressed protein coding genes (RPKM>=1) in hESCs. For analysis in endoderm differentiation, the background was defined as all expressed protein-coding genes (RPKM>=1) in endoderm differentiation (GSE75297) (Daneshvar et al., 2016).

### Statistical analyses of large-scale data

The p-values for large-scale comparison were calculated by Wilcoxon–Mann–Whitney test unless otherwise stated.

## Acknowledgments

We would like to thank Ross Tomaino and the Taplin Mass/spec core at Harvard Medical School, the MGH nextGen Sequencing Core, the MGH PMB Microscopy Core Facility, and the HSCI-CRM Flow Cytometry Core Facility at MGH. We thank Tomoe Kitao Ando (Gifu University) for providing consultation in the optimization of the RNA pull-down protocol. We are thankful for the continuous help and collaboration from Marc Beal, Josh Ryu, and Ron Cook of Optical Biosystems for their assistance with RNA detection and high-throughput microscopy. This work was supported by NIH/NICHD grant R01HD09277302 to A.C.M.

---

**Figure S1.**
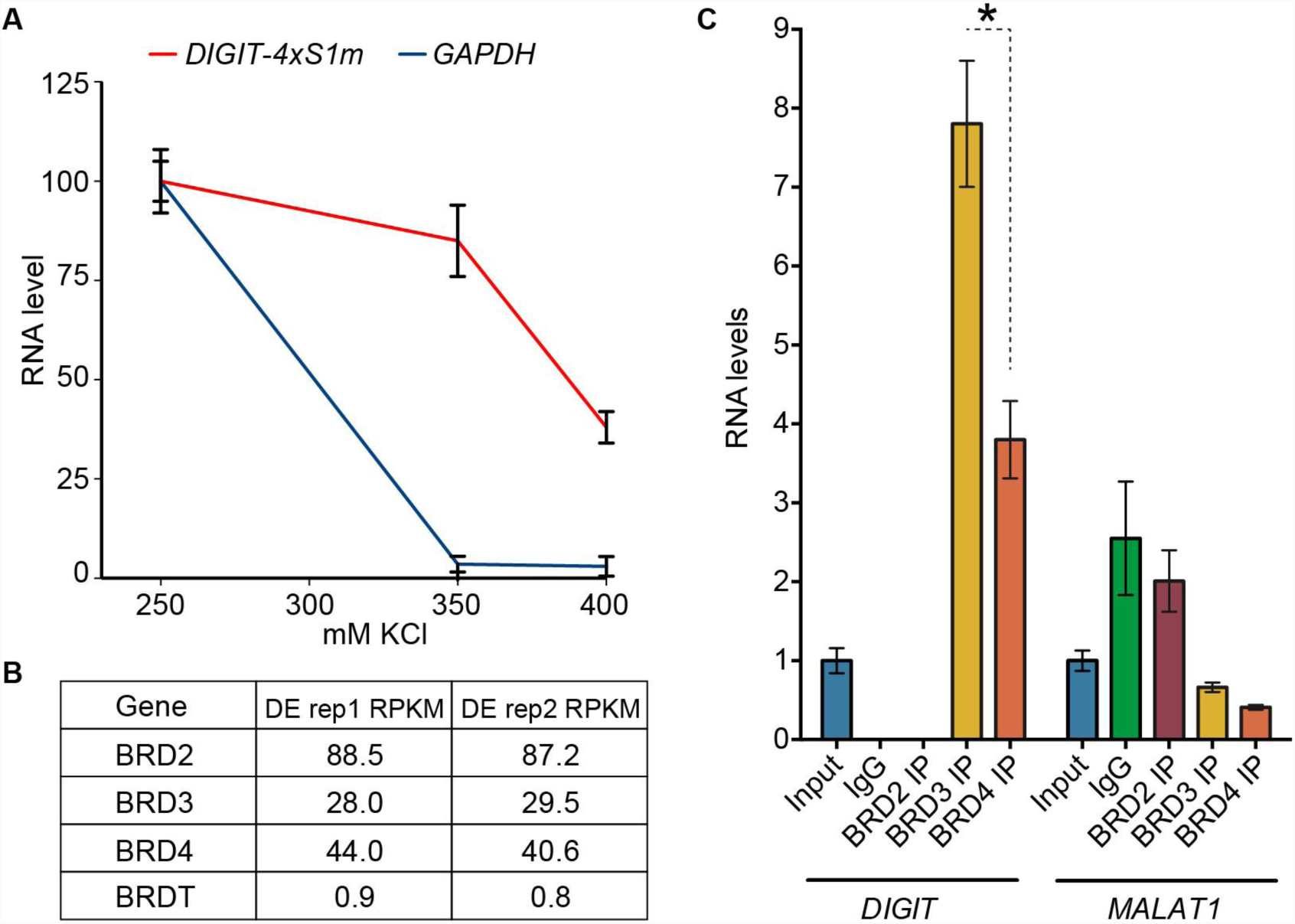
Optimization of washes for RNA pull-down, expression of BRD family members in endoderm, and binding of *DIGIT* to BRD proteins. **(A)** Recovery of *DIGIT-4S1m* versus endogenous *GAPDH* mRNA with increasing concentrations of KCl. The amount of RNA recovered with 250 mM KCl wash conditions is set to 100 for each RNA species. Error bars represent standard deviation of three replicates. **(B)** The table shows normalized RPKM values for expression of BRD2, BRD3, and BRD4 with endoderm differentiation (Daneshvar et al., 2016). **(C)** qRT-PCR shows the enrichment *DIGIT* (left) and *MALAT1* (right) following immunoprecipitation of BRD2, BRD3, and BRD4 proteins on day 4 of endoderm differentiation. IgG is used as a non-specific control. Enrichment is quantified relative to *GAPDH*. * indicates p <0.05.

**Figure S2.**
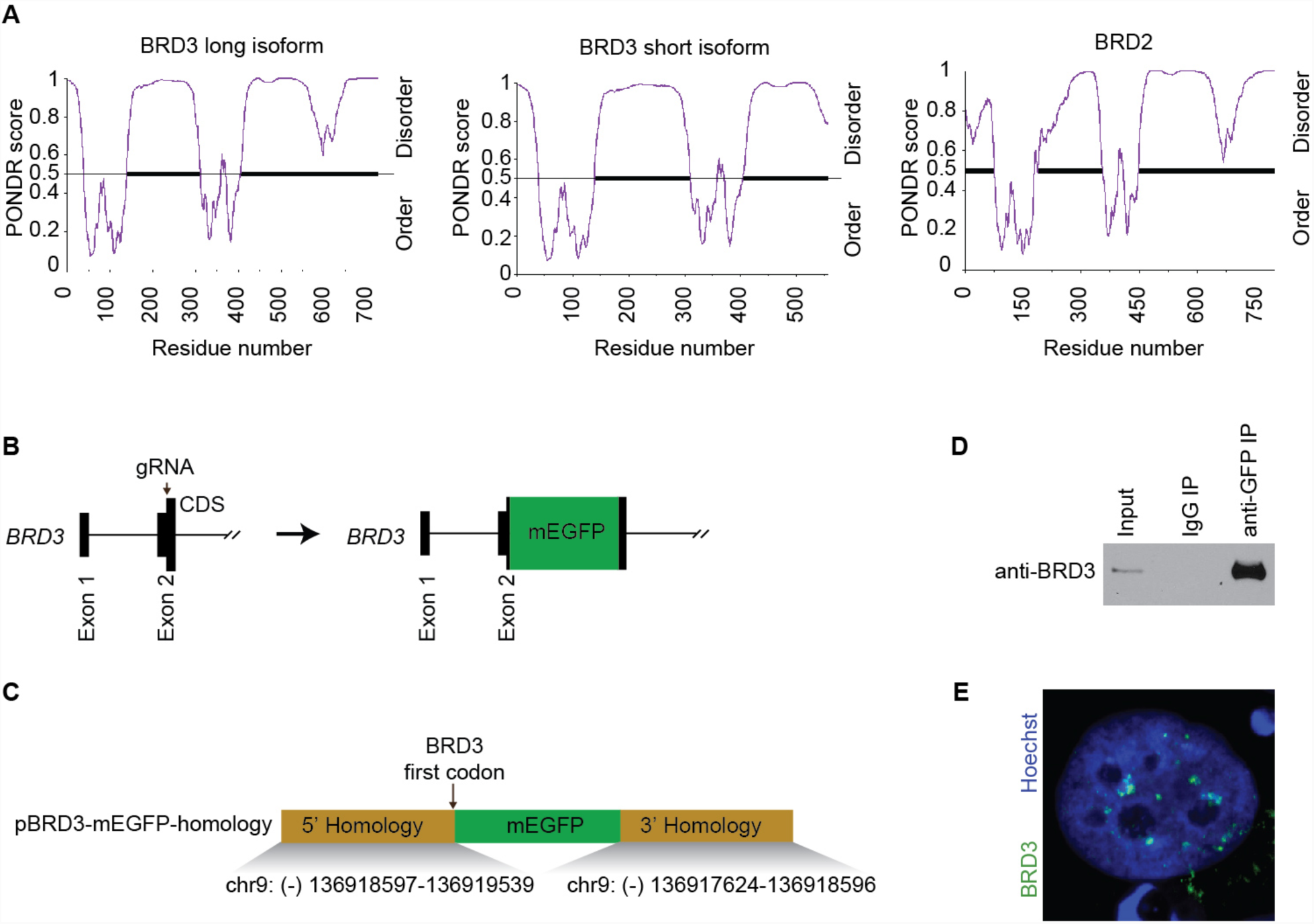
Prediction of ordered/disordered protein structures, strategy for tagging the endogenous BRD3 with mEGFP, and formation of BRD3 puncta in hESCs. **(A)** PONDR VSL2 plots showing the ordered and disordered regions of the long (left) and short (middle) isoforms of BRD3, and BRD2 (right). **(B)** Creation of mEGFP-BRD3 fusion protein. The cDNA encoding mEGFP was inserted at the N-terminus of *BRD3*. This terminus is shared by the long and short isoforms of BRD3. The site targeted by gRNA is indicated with an arrow. **(C)** Design of the homology vector for insertion of mEGFP. The genomic locations of homology arms are indicated. Arrow indicates that mEGFP was inserted after the start codon for *BRD3*. **(D)** Immunoblot using anti-BRD3 antibody confirms the generation of mEGFP-BRD3 fused protein. hESCs were lysed and IP was performed using IgG isotype control and an antibody recognizing GFP. Total cell lysates (input) and the IPs were then probed with an anti-BRD3 antibody. **(E)** Immunofluorescence of an hESC cell showing BRD3 (green) in the nucleus (blue) of the hESCs.

**Figure S3.**
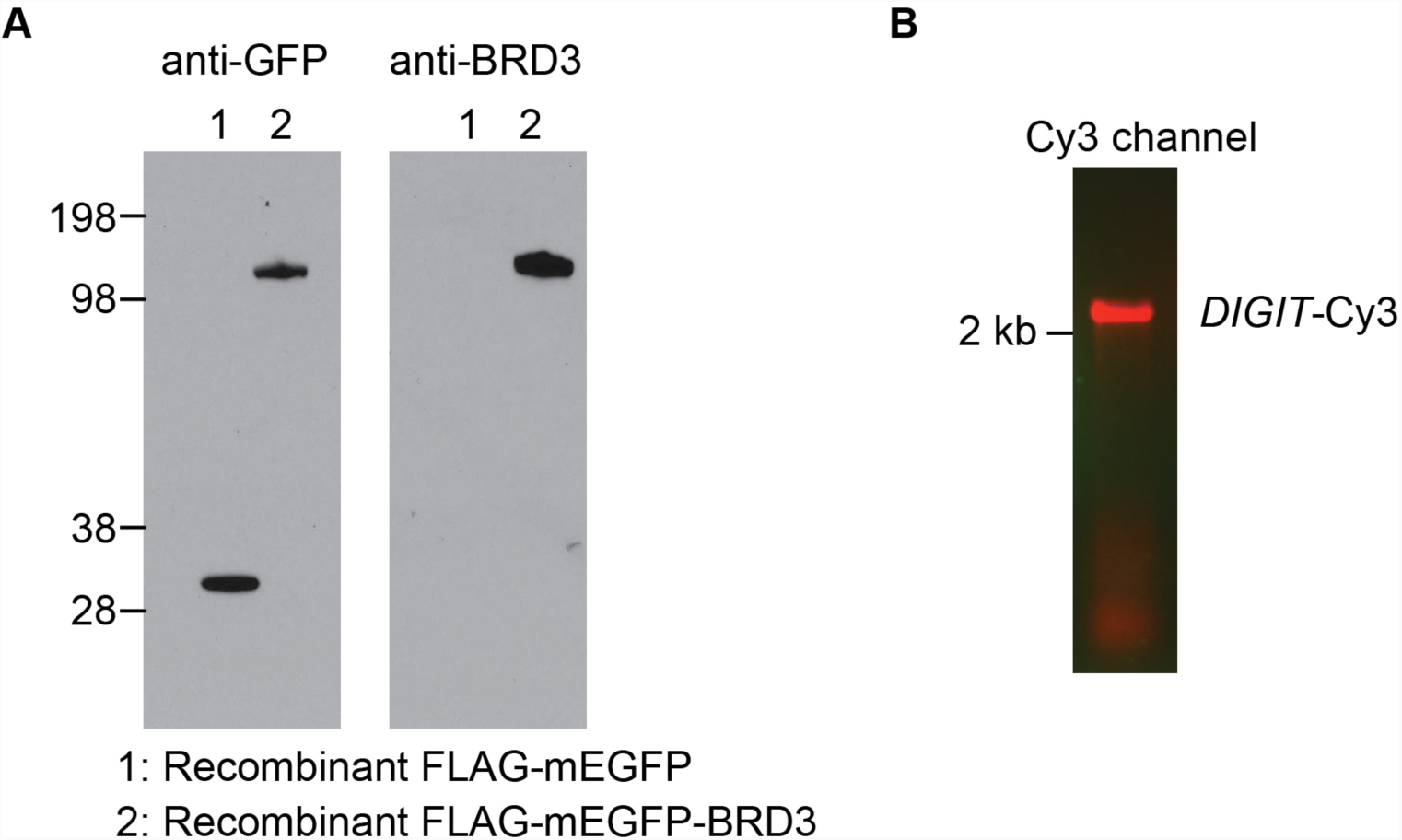
Production of recombinant BRD3 and *in vitro* transcribed and labeled *DIGIT*. **(A)** Immunoblot was performed to detect recombinant FLAG-mEGFP-BRD3 and FLAG-mEGFP using an anti-GFP antibody (left) and an anti-BRD3 antibody (right). **(B)** Fluorescent detection of the Cy3-labeled *DIGIT* on a 1% bleach-agarose gel.

**Figure S4.**
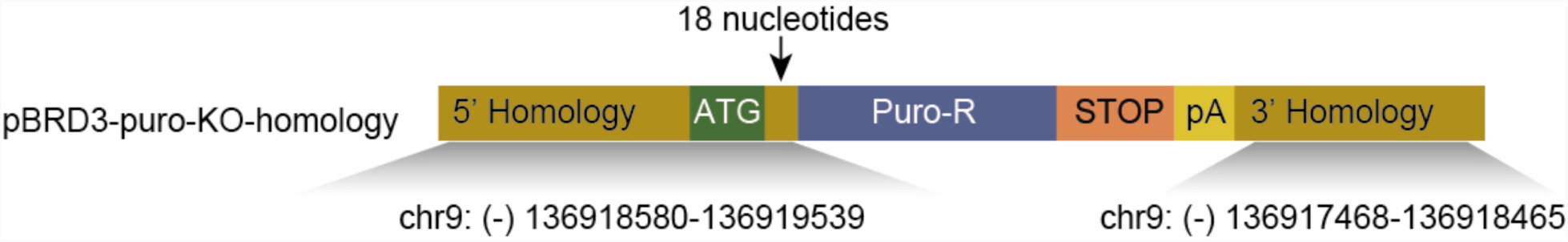
Homology construct for targeting BRD3. Map of the homology construct for insertion of a puromycin resistance cassette and a stop codon cassette downstream of the sixth codon of the gene encoding BRD3.

**Figure S5.**
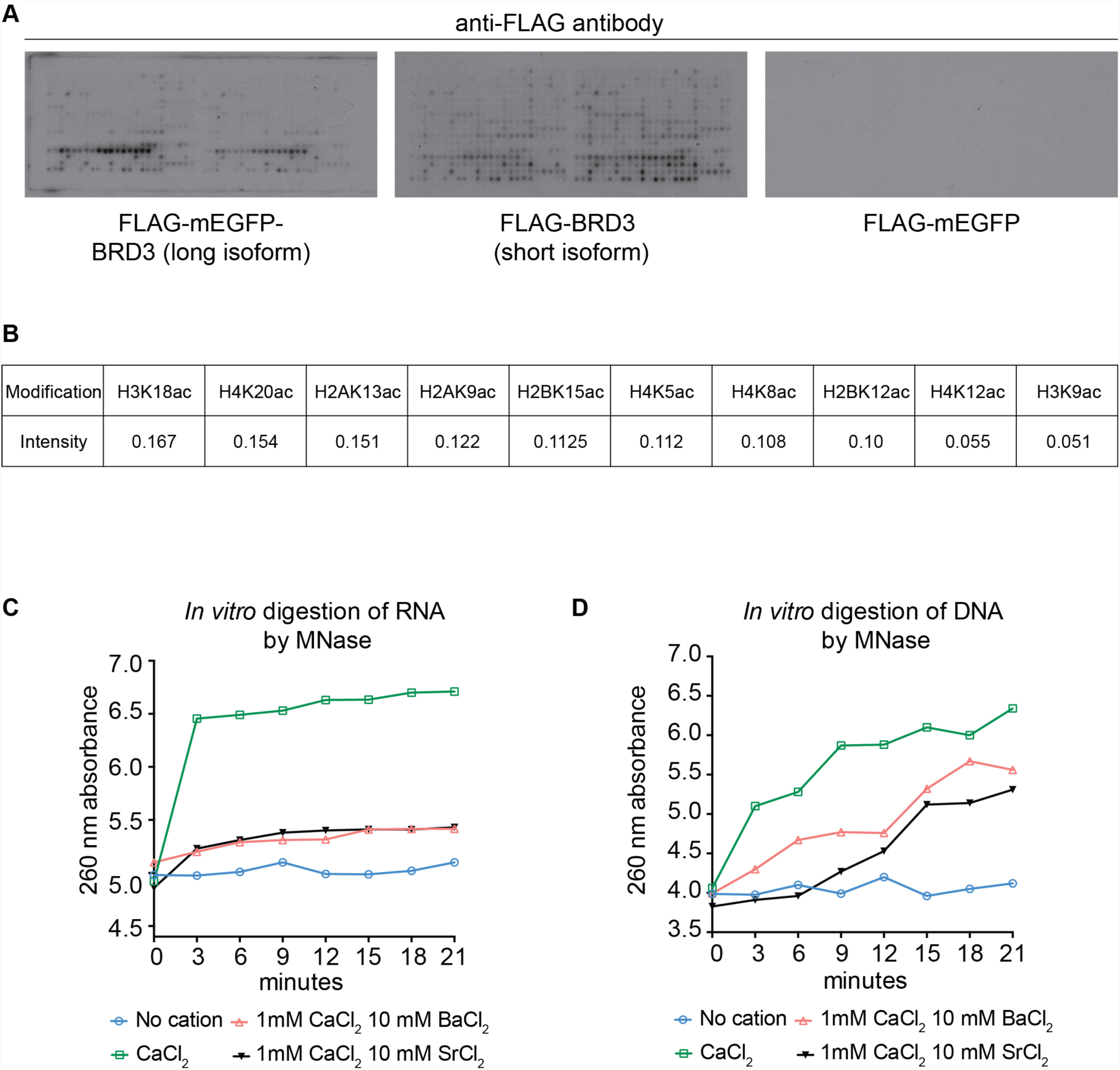
Binding of BRD3 to modified histones and activity of MNase in presence of of Ba^+2^ and Sr^+2^ cations. **(A)** Peptide arrays show the binding of recombinant BRD3 to histone modifications. **(B)** Spot intensity of BRD3 (long isoform) binding to the top ten histone modification as quantified by image processing software (see materials and methods). **(C)** RNase activity of MNase in presence of Ba^+2^ and Sr^+2^ cations. **(D)** DNase activity of MNase in presence of Ba^+2^ and Sr^+2^ cations.

**Figure S6.**
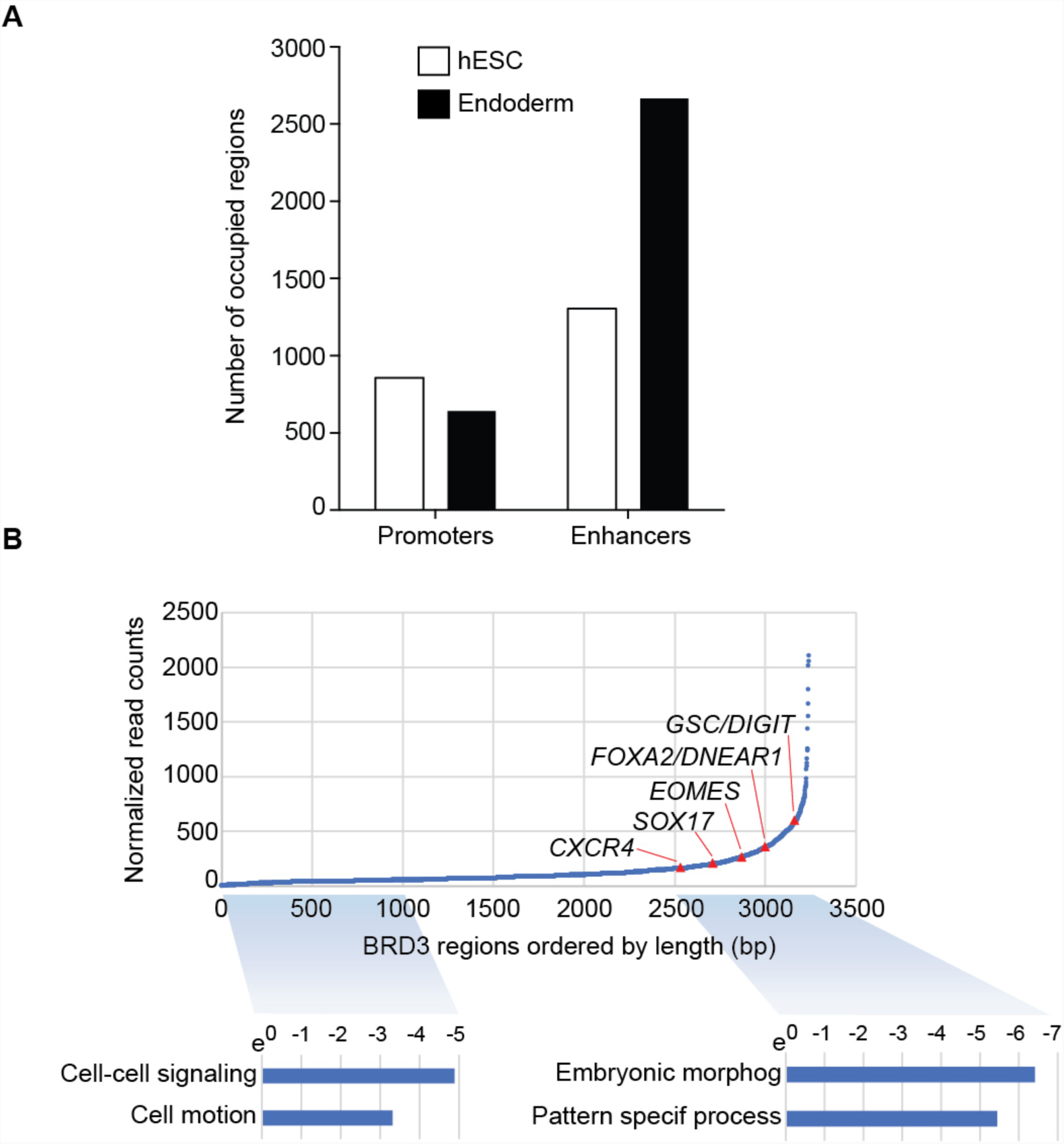
BRD3 occupies endoderm genes and enhancers. **(A)** The number of BRD3 regions (y-axis) that contain promoters and enhancers are shown for hESCs (white) and endoderm cells (black). BRD3 regions are defined as containing promoters if the region is located within 2kb of a transcription start site (TSS). BRD3 regions are defined as containing enhancers if the region overlaps with enhancers defined by H3K27ac. Regions that contain both promoters and enhancers are counted in both categories. **(B)** Regions of BRD3 occupancy (3274) from cells on day 4 of endoderm differentiation were ordered by increasing length (x-axis). The normalized read counts for each occupied region are shown on the y-axis. Red triangles indicate the location of endoderm genes. GO analysis was performed for the genes associated with regions of BRD3 occupancy (bottom). The 1000 BRD3 regions with the smallest length (blue shading, left) were associated with 373 genes. The regions of BRD3 occupancy with the largest length (starting with region 2500, blue shading, right) were associated with 348 genes. The top two categories from GO analysis are shown for each set of genes.

**Figure S7.**
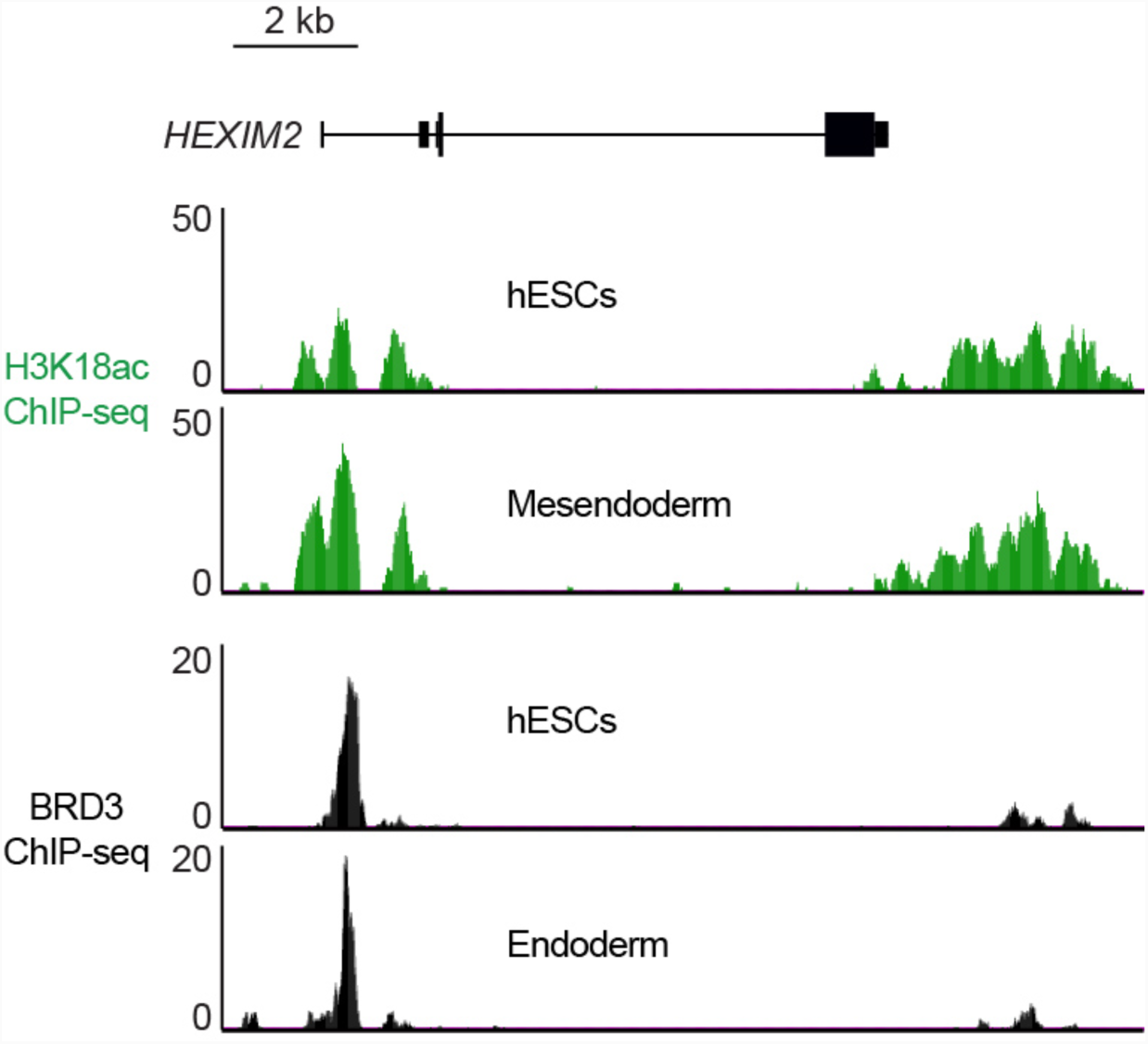
BRD3 occupies *HEXIM2* locus in hESCs and endoderm. ChIP-seq data shows H3K18ac (top, green) and BRD3 occupancy (bottom, black) at *HEXIM2* in hESCs and endoderm / mesendoderm cells.

